# Genital tract microbiome dynamics are associated with time of *Chlamydia* infection

**DOI:** 10.1101/2022.07.18.500533

**Authors:** Lihong Zhao, Stephanie R. Lundy, Francis O. Eko, Joeseph U. Igietseme, Yusuf O. Omosun

## Abstract

**Background:** We have previously shown that the time of *Chlamydia* infection was crucial in determining the chlamydial infectivity and pathogenesis. This study aims to determine whether the time of *Chlamydia* infection affects the genital tract microbiome. This study analyzed mice vaginal, uterine, and ovary/oviduct microbiome with and without *Chlamydia* infection. The mice were infected with *Chlamydia* at either 10:00 am (ZT3) or 10:00 pm (ZT15).

**Results:** The results showed that mice infected at ZT3 had higher *Chlamydia* infectivity than those infected at ZT15. There was more variation in the compositional complexity of the vaginal microbiome (alpha diversity) of mice infected at ZT3 than those mice infected at ZT15 throughout the infection within each treatment group, with both Shannon and Simpson diversity index values decreased over time. The analysis of samples collected four weeks post-infection showed that there were significant taxonomical differences (beta diversity) between different parts of the genital tract—vagina, uterus, and ovary/oviduct—and this difference was associated with the time of infection. *Firmicutes* and *Proteobacteria* were the most abundant phyla within the microbiome in all three genital tract regions for all the samples collected during this experiment. Additionally, *Firmicutes* was the dominant phylum in the uterine microbiome of ZT3 *Chlamydia* infected mice.

**Conclusion:** The results show that the time of infection is associated with the microbial dynamics in the genital tract. And this association is more robust in the upper genital tract than in the vagina. This result implies that more emphasis should be placed on understanding the changes in the microbial dynamics of the upper genital tract over the course of infection.

## 1 Background

*Chlamydia trachomatis* (*C. trachomatis*), a gram-negative, obligate intracellular bacteria, causes the disease *Chlamydia. Chlamydia* causes trachoma, respiratory disease, and sexually transmitted infection [1, 2]. There are an estimated 130 million *Chlamydia* cases worldwide and an estimated 3 million new cases annually in the United States, with many unreported cases due to their asymptomatic nature [1, 2]. Sexually active individuals are at risk of having *Chlamydia* infection; two-thirds of new infections occur among young women aged 15 to 24 [1, 2]. After infection, *C. trachomatis* migrates into the upper genital tract [1, 2, 3], where it infects the cervix and urethral columnar epithelial cells, causing uncomplicated cervicitis and urethritis. This infection leads to symptomatic and asymptomatic pelvic inflammatory disease (PID) experienced by 2-5% of women with untreated *Chlamydia* infection. PID can cause irreversible damage to the uterus, fallopian tubes, and surrounding tissues leading to pelvic pain, tubal factor infertility, and potentially fatal ectopic pregnancy [1, 2]. Some symptoms include urinary tract infection, vaginal discharge, and bleeding [1, 2, 4]. While *Chlamydia* infection can lead to these devastating pathologies, we see varying levels in the severity of the disease outcomes in women exposed to *Chlamydia*. Exposure to the bacteria does not always lead to an infection, and it does not always result in severe disease complications. This phenomenon suggests that several factors, including host immune response, pathogen virulence, the time of infection, and the vaginal microbiome, may impact the susceptibility and varying disease outcomes observed in women infected with *Chlamydia*.

Several physiological, cellular, and behavioral processes exhibit a daily circadian rhythm without external cues. These rhythms are controlled by the body’s internal clock and describe the endogenous oscillation in an organism observed in approximate association with the Earth’s daily rotation [5, 6, 7]. Circadian rhythms managed by the light/dark cycle allow organisms to anticipate environmental changes. In mammals, circadian rhythms are regulated by the molecular clock of the hypothalamus suprachiasmatic nucleus and circadian clocks in most peripheral tissues [8]. Studies have reported circadian rhythms controlling host immune response, reproduction, antibacterial host defenses, sepsis, inflammation, and cell proliferation [7, 9, 10, 11]. As understanding of the role of circadian rhythms on infection has grown, studies conducted on pathogens such as human herpesvirus 2, influenza, *Salmonella typhimurium*, and *Chlamydia* not only suggest that circadian rhythms influence the progression of the disease, but that the time of infection seems to play an essential role in overall disease outcomes [12, 13, 14].

A lot of studies have been performed on the human vaginal microbiome, which has high diversity, and is dominated by *Lactobacilli* which helps maintain health and prevent disease [15, 16, 17]. The microbiota changes quickly to a state linked with a disease and is associated with the metabolome [18]. Most microbiome studies have focused on the vaginal and rectal microbiome, or the endocervix [17, 19, 20]. It has been reported that other parts of the female reproductive system that were once assumed to be immune-privileged also have microbes living there [21, 22]. *Chlamydia* has been associated with changes in the vaginal microbiome, associated with PID [23, 24, 25]. Transcriptomic analysis shows crosstalk between the host and microbiota involving histone deacetylase-controlled pathways [26]. Some studies have shown that differences in the microbial taxa abundance rather than the bacterial taxa are essential in *Chlamydia* infection [27]. The microbiome is also been associated with selected inflammatory mediators [28].

Studies have shown that chlamydia pathogenesis is partly regulated through circadian rhythms [14, 29]. This understanding was prompted by noticing that the time of infection was important in delineating chlamydia burden and pathogenesis [14]. In addition, the disruption of host circadian rhythms was associated with worse pathogenic outcomes [29]. While the host circadian clock can regulate some viral and bacterial infections, including *Chlamydia* [12, 13, 14], we do not fully understand the role of host circadian rhythms on the pathogenesis of genital *Chlamydia* infection and the possible role of the entire genital tract microbiome, not only the vagina, in developing the adverse pathology associated with chlamydial infection and pathogenesis. We hypothesize that chlamydia infection and the eventual chlamydial burden will depend on the genital tract microbial communities at the time of infection. The reverse will also be the case with the distribution and richness of the genital tract microbial community depending on the time of chlamydia infection. These microbial population-based changes related to infection time could help us better understand the processes involved in *Chlamydia* pathogenesis, especially the mechanism leading to infertility in women.

## 2 Methods

### 2.1 Animals

Female C57BL/6J mice (Jackson Laboratory, Bar Harbor, MA) received at six weeks old were housed under normal light: dark cycle conditions of 12 hours light: 12 hours dark (12:12LD). The light intensity during the light phase in the room was 906 Lux.

### 2.2 *Chlamydia muridarum* Stock

*Chlamydia muridarum niggs* (*C. muridarum*) stocks (Centers for Disease Control, Atlanta, GA) were diluted in sterile sucrose phosphate glutamate transport media to a final concentration of 1 × 105 Infectious Units (IFUs).

### 2.3 *C. muridarum* Infectivity Assay

All mice were subcutaneously injected between 10:00 AM and 12:00 PM with 2.5 mg/mouse Depo Provera, medroxyprogesterone acetate (Pfizer, New York, NY) in sterile Phosphate Buffer Saline to synchronize the estrous cycle. Mice were intravaginally infected seven days later, with 1 × 105 *C. muridarum* at 10:00 AM Zeitgeber time (ZT) 3, early rest period) or 10:00 PM (ZT15, early active period), which can be interpreted as three hours after the lights were turned on or off in the room (7:00 AM lights on, 7:00 PM lights off), respectively. The mice were anesthetized using isoflurane during the process of infection. Infected mice were swabbed every three days for 27 days, and the bacteria were isolated and cultured to track the progression and clearance of the infection.

### 2.4 Swab and Tissue Collection for Microbiome Analysis

Uninfected and infected mice were swabbed before infection and once a week for four weeks post-infection to determine the microbial community composition at each time point. Note that the first set of swab samples (before infection) were samples collected the same day the mice were (sham and chlamydia) infected, we denote them as week 0 samples and consider week 0 as the baseline. The second set of swab samples were collected one week after infection, we denote them as week 1 samples. Thirty days post-infection, all mice were sacrificed between 10:00 AM and 12:00 PM, and the genital tract was removed and divided into two groups: the ovary and oviduct, and the uterus. Euthanasia was performed using carbon dioxide and cervical dislocation. The swabs and genital tracts were stored at −20 °C until DNA was extracted from the samples.

### 2.5 DNA Extraction, Amplification, Metagenomic Sequencing

DNA from the vaginal swabs and genital tract tissues (190 samples in total) were extracted using the QIAamp DNA Micro Kit (Qiagen), following standard procedure, and quantified using nanodrop. Genital tract tissues were dissociated using Collagenase/Dispase Blend I (Millipore) and incubated in a water bath at 37 °C before following the QIAamp DNA Micro Kit protocol. Hypervariable regions V3-V4 of the small subunit ribosomal RNA (16s rRNA) genes were amplified using eubacterial primers. Forward primer 5’ TCGTCGGCAGCGTCAGATGTGTATAAGAGACAGC-CTACGGGNGGCWGCAG and reverse primer 5’ GTCTCGTGGGCTCGGAGATGTGTATAAGAGACAGGACTACHVGGGTATCTAATCC were fused with Illumina overhang adapters and specific dual index barcodes to allow multiple samples to be analyzed on a single picotiter plate. The pooled DNA was amplified with PCR (Amplicon PCR Reverse Primer, Amplicon PCR Forward Primer, 2x KAPA HiFi HotStart Ready Mix) under the following amplification protocol conditions: 95 °C for 3 minutes, 25 cycles of 95 °C for 30 seconds, 55 °C for 30 seconds, 72 °C for 30 seconds, 72 °C for 5 minutes. The PCR product cleanup was performed by using AMPure XP beads before the final library was quantified. Purified amplicons were processed according to Illumina standard protocol and paired 300 base pair sequencing was conducted on the MiSeq Illumina platform with the reagent kit V3 at the Georgia Genomics and Bioinformatics Core at the University of Georgia.

### 2.6 Sequence Data Processing

All analyses were performed using R version 4.1.0 with DADA2 version 1.20.0. Sequence reads were first pre-processed to trim off the primer sequence and then processed through DADA2 pipeline to identify amplicon sequence variants (ASVs) with chimeras being removed. These ASVs were classified to the genus level using the RDP naive Bayesian classifier in combination with the SILVA reference database version 138. Phylogenetic tree was constructed using DECIPHER version 2.20.0 and phangorn version 2.7.0 with sequences extracted from DADA2 output. For our analyses, the phylogenetic tree was rooted by the longest branch terminating in a tip.

Singletons were removed for all downstream analyses. A total of 3165 ASVs were identified in 190 samples, and these data were used to measure alpha diversity. The smallest number of reads per sample is 5982. For subsequent analyses, we further removed ASVs with ambiguous phylum annotation and ASVs belonging to a phylum with total prevalence number less than 4. This led to a total of 2870 ASVs in 190 samples.

### 2.7 Statistical Analysis and Data Visualization

Two–way analysis of variance (ANOVA) was used to determine the difference in infectivity between the treatment groups. We did a Tukey post hoc test (for multiple comparisons by comparing the mean of each group to the mean of every other group) after the two–way ANOVA to determine the actual statistical relationship between the treatment groups. Statistical significance was determined at *p* < 0.05. GraphPad Prism (La Jolla, CA) was used to performed the above mentioned analyses. All the following analyses were performed using R version 4.1.0. Seed 42 was used for the Marsaglia-Multicarry random number generator for the results presented in this paper. We performed pairwise Kruskal-Wallis rank-sum test to evaluate significant differences in alpha diversity index among groups for samples collected per week, and to evaluate significant differences in alpha diversity index between genital tract regions (GTRs) for samples collected in week 4 (i.e., 4 weeks post-infection). A significant Kruskal-Wallis test was followed with post-hoc analysis of multiple comparisons using a Dunn’s test with Bonferroni adjustment. For vaginal samples, we conducted pairwise comparisons using the Wilcoxon rank-sum test to evaluate significant differences in alpha diversity index between samples collected in different time within each group.

To test whether the overall microbial community differs by variables of interest (group, week, GTR), we performed permutational multivariate analysis of variance (PERMANOVA) with vegan version 2.5.7, using three distance measures: Bray-Curtis measure of dissimilarity, UniFrac, and weighted UniFrac (W-UniFrac). *P* values for the test statistic (pseudo-*F*) were based on 999 permutations. We also performed a permutation-based test of multivariate homogeneity of groups dispersions (betadisper) to identify pairs that differ significantly in dispersion (variance) for each PERMANOVA test. To visualize differences in beta diversity according to variables of interest (group and week or group and GTR), we applied principal coordinate analysis (PCoA) to Bray-Curtis measure of dissimilarity, UniFrac, and weighted UniFrac (W-UniFrac), respectively.

For samples collected four weeks after *Chlamydia* infection, we further performed Canonical Correspondence Analysis (CCA) to identify candidate indicator ASVs that respond to the variable of interest (GTR within each group or group within each GTR). A Venn diagram was generated to show the shared and unique ASVs among GTRs, based on the occurrence of ASVs in samples collected four weeks after infection regardless of their abundance.

## 3 Results

To evaluate whether the time of day of pathogen exposure impact the genital tract microbiome in *Chlamydia* infection, we compare the alpha diversity metrics within individual samples and beta diversity metrics between pairs of samples, and conduct PERMANOVA and betadisper analyses on vaginal samples collected before and post-infection as well as samples collected in different genital tract region four weeks post-infection. We observed some differences in both alpha and beta diversity metrics of vaginal samples collected over the course of infection within each treatment group, which suggest the time of infection is important in *Chlamydia* infection. The CCA analyses carried out on samples collected four weeks post-infection indicate that the effect of time of pathogen exposure on *Chlamydia* infection is stronger in the upper genital tract than in the vagina. We were able to identify several candidate ASVs that may be helpful in deciphering this association.

### 3.1 Time of Day of Pathogen Exposure is Important in Chlamydial Infectivity

Chlamydia burden was determined by measuring the shedding of chlamydiae into the cervicovaginal vault. We have previously shown that mice infected with *C. muridarum* at ZT3 had significantly higher bacteria burden than the mice infected at ZT15, 12-24 days post-infection [14]. The *Chlamydia* burden in this study is similar to our previous study with mice infected at ZT3 having higher *Chlamydia* burden than mice infected at ZT15, although not statistically significant in this case (Fig. 1). The results reiterate that the time of day when mice were exposed to the pathogen is essential.

**Figure 1:**
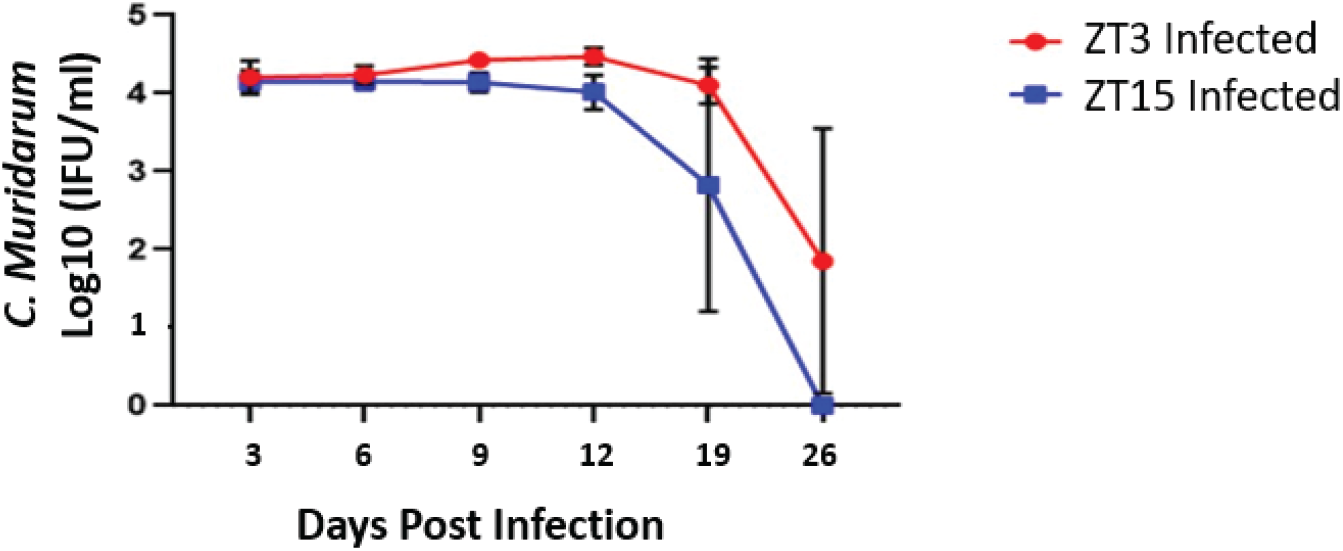
Chlamydia infectivity of mice housed under 12:12LD. Mice (*n* = 10 per group) were infected with *C. muridarum* at either ZT3 or ZT15. Mice infected at ZT3 had a higher *Chlamydia* burden compared to mice infected at ZT15. Data were analyzed using a two-way analysis of variance (ANOVA). The differences in infectivity between ZT3 and ZT15 infected mice were not statistically significant.

### 3.2 Vaginal Samples

Vaginal swab samples were collected at five time points: week 0, week 1, week 2, week 3, week 4, and week 5. The pre-infection samples collected in week 0 are our baseline samples.

#### 3.2.1 Alpha diversity of vaginal microbiome differ with time of infection

Alpha diversity measures the compositional complexity within one sample. Many alpha diversity indices exist and each reflects different aspects of community heterogeneity. Two diversity metrics were used in our analysis: Shannon diversity index and Simpson diversity index. Both metrics account for richness (the number of species present) and evenness; the diversity index value increases as the richness or evenness of the community increases, with Simpson diversity index being less sensitive to rare species than Shannon diversity index. The distribution of both Shannon diversity index and Simpson diversity index of vaginal samples from mice infected at ZT3 (ZT3_I) and ZT15 (ZT15_I) along with their corresponding control groups (ZT3_C and ZT15_C) changes with time, as shown in Figs. 2 and A.2. However, these changes in both diversity indices were not statistically significant (*p* > 0.05, Kruskal-Wallis rank-sum test).

**Figure 2:**
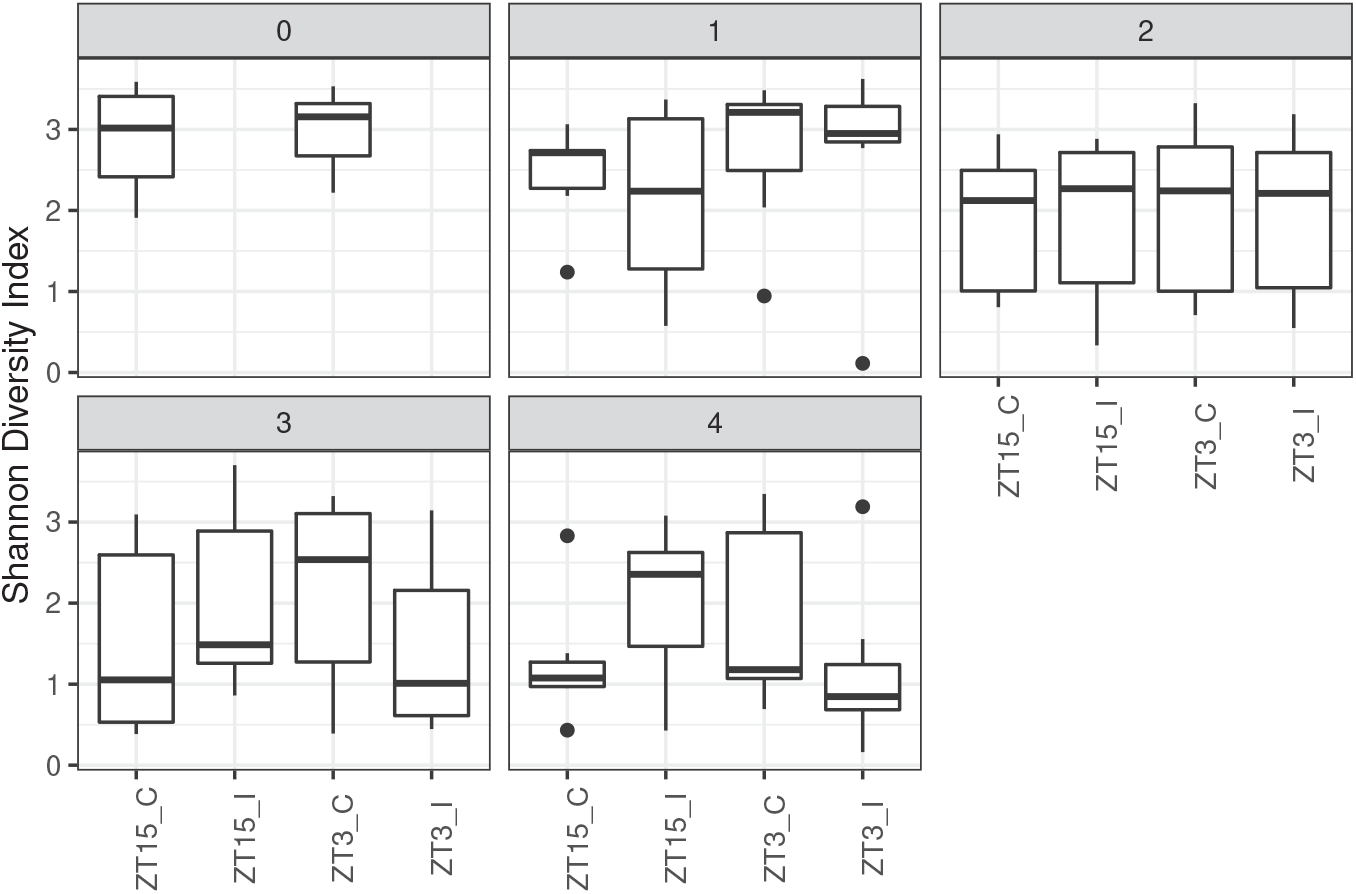
Boxplots of Shannon Diversity Index for vaginal samples by group (ZT15_C, ZT15_I, ZT3_C, and ZT3_I) per week. The thick line within the box represents the median. The lower and upper hinges correspond to the first and third quartiles (the 25th and 75th percentiles). The upper and lower whiskers extend from the hinge to the largest or smallest value no further than 1.5×IQR (inter-quartile range, the distance between the first and third quartiles), respectively. Individual dots represent “outlying” points which were not removed from the analysis.

In the sham control groups (ZT3_C and ZT15_C), the values of both Shannon and Simpson diversity indices of vaginal samples collected in week 2, 3, and 4 were significantly lower than the ones collected in week 0, i.e., the pre-infection baseline (*p* ⩽ 0.01, Wilcoxon rank sum test), as shown in Fig. A.1 in Supplementary Material. Additionally, as shown in Fig. A.1, for the sham control groups, the values of both diversity indices of vaginal samples collected four weeks after infection were significantly lower than the ones collected one week after infection (*p* ⩽ 0.05, Wilcoxon rank sum test); while for infected group (ZT15_I and ZT3_I), the difference between vaginal samples collected four weeks after infection and the ones collected one week after infection is only significant with Simpson diversity index (*p* ⩽ 0.05, Wilcoxon rank sum test). When the comparisons were performed within each treatment group (ZT15_C, ZT15_I, ZT3_C, or ZT3_I), ZT15_C samples collected three and four weeks post-infection had significant lower Shannon diversity index value compared to samples collected in week 0, the baseline before sham infection (*p* ⩽ 0.05, Wilcoxon rank sum test). However, the differences of Simpson diversity index for mice infected at different times were no longer statistically significant (*p* > 0.05, Wilcoxon rank sum test), as shown in Fig. 3. It should be noted that the values of both diversity indices for vaginal samples of mice infected at ZT3 were decreasing over time, while less changes observed in mice infected at ZT15 (Supplementary Fig. A.3).

**Figure 3:**
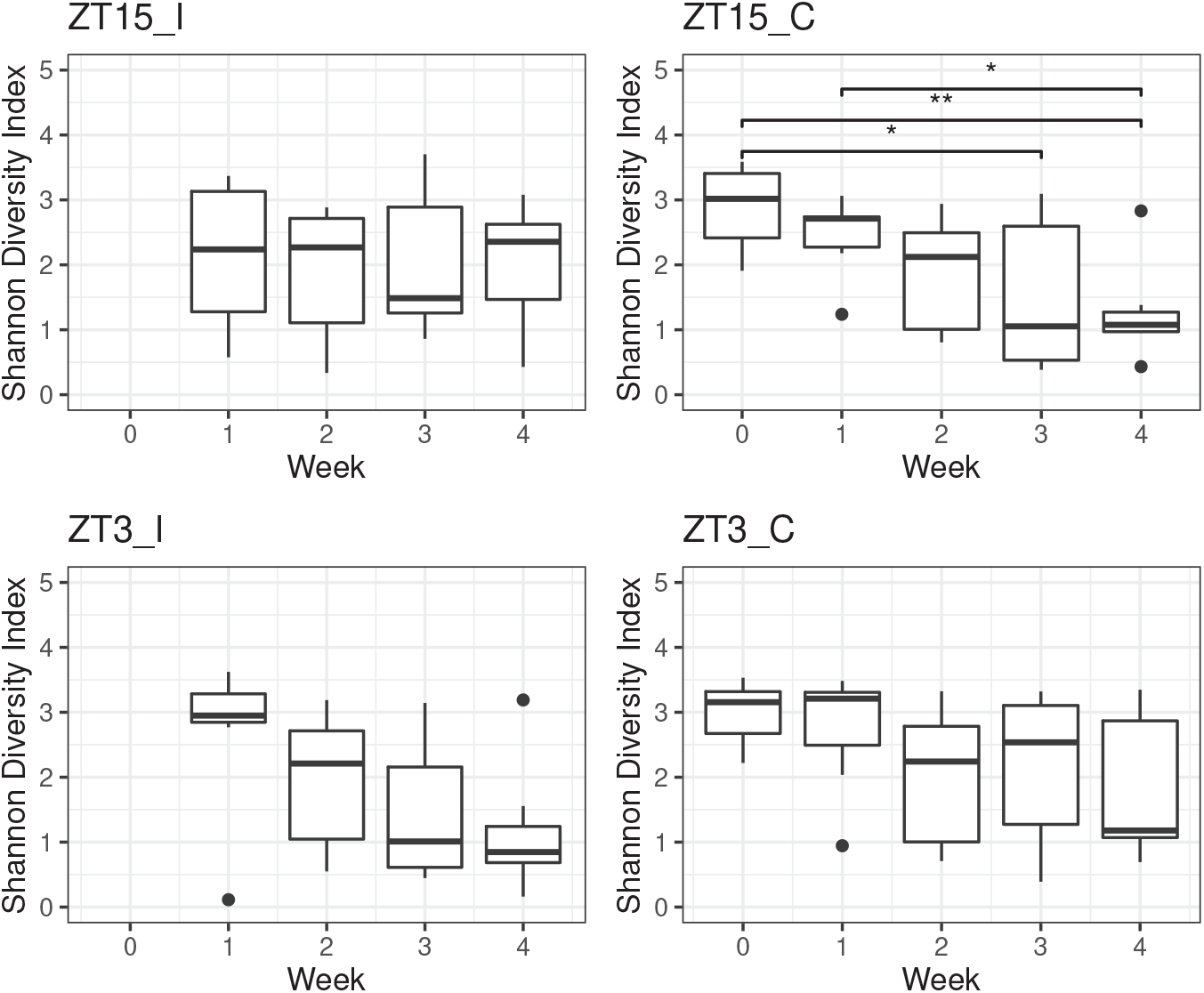
Boxplots of Shannon Diversity Index for vaginal samples per group (ZT15_C, ZT15_I, ZT3_C, and ZT3_I) over time. Statistical significance is indicated above the brackets: *,*p* ⩽ 0.05; **,*p* ⩽ 0.01.

#### 3.2.2 Beta diversity of vaginal microbiome differ with time of infection

Beta diversity measures the taxonomical similarity or dissimilarity between pairs of samples. Three metrics were used to measure beta-diversity: Bray-Curtis measure of dissimilarity, UniFrac, and W-UniFrac. Bray-Curtis measure takes into account both the presence/absence and abundance of ASVs but not phylogeny. Both UniFrac and W-UniFrac takes into account phylogenetic relationships of ASVs present in the microbiota, W-UniFrac weights the branches of a phylogenetic tree based on the abundances of ASVs and is less sensitive to low abundance ASVs and those ASVs that are very phylogenetically distant compared to UniFrac (which only accounts for the presence/absence of ASVs and not abundance).

For all vaginal samples, we observed that the period after infection in weeks, i.e., the independent variable week, had a significant impact on the vaginal microbiota composition with all three distance metrics we tested. However, the significance with Bray-Curtis and W-UniFrac might be partially due to the differences in dispersion between samples collected in week 0 and week 2; while the significance with Unifrac might be partially due to the differences in dispersion between other pairs of samples, as shown in Table 1 and Fig. 4. We also observed that the vaginal microbiota composition differ among Groups (ZT15_C, ZT15_I, ZT3_C, and ZT3_I) with Bray-Curtis and UniFrac but not W-UniFrac. However, this might be due to the differences in dispersion between samples in ZT15 C and ZT3_C (Bray-Curtis and UniFrac) as well as ZT15_I and ZT3_C (UniFrac), as shown in Table 1 and Fig. 5.

**Table 1:**
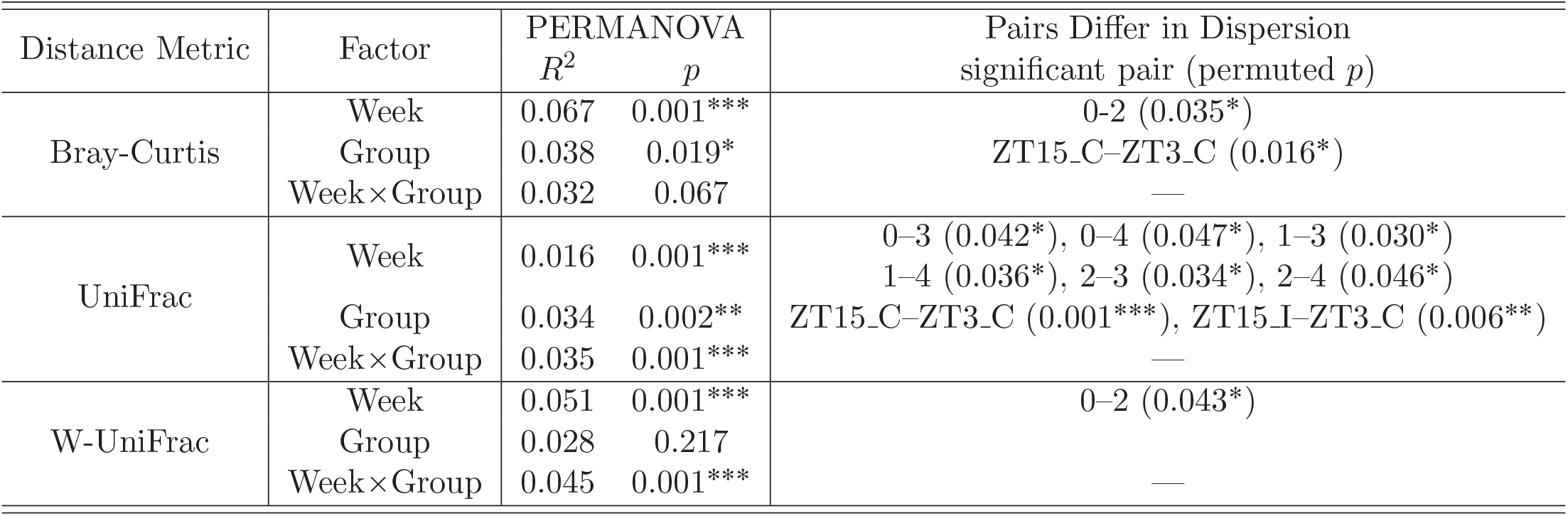
PERMANOVA and betadisper results for all vaginal samples. Asterisk denotes statistically significant difference (*,*p* ⩽ 0.05; **,*p* ⩽ 0.01; ***,*p* ⩽ 0.001).

**Figure 4:**
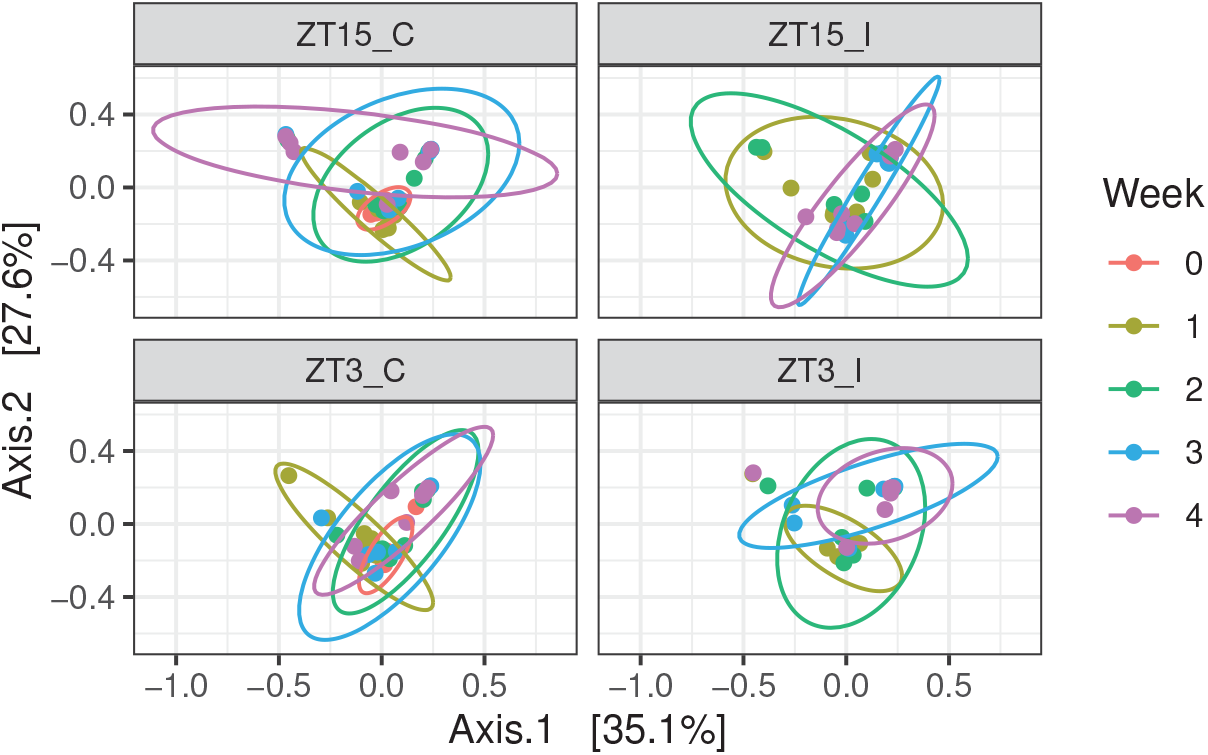
PCoA ordination plots for beta diversity metrics before (week 0) and post-infection (week 1, 2, 3, 4) by treatment groups, showing W-UniFrac, for vaginal samples. Each dot represents the microbial community of a sample, the color of the dots corresponds to the week the sample was collected. The first axis explains 35.1% of the variability and the second axis explains 27.6% of the variability in the data of vaginal samples.

**Figure 5:**
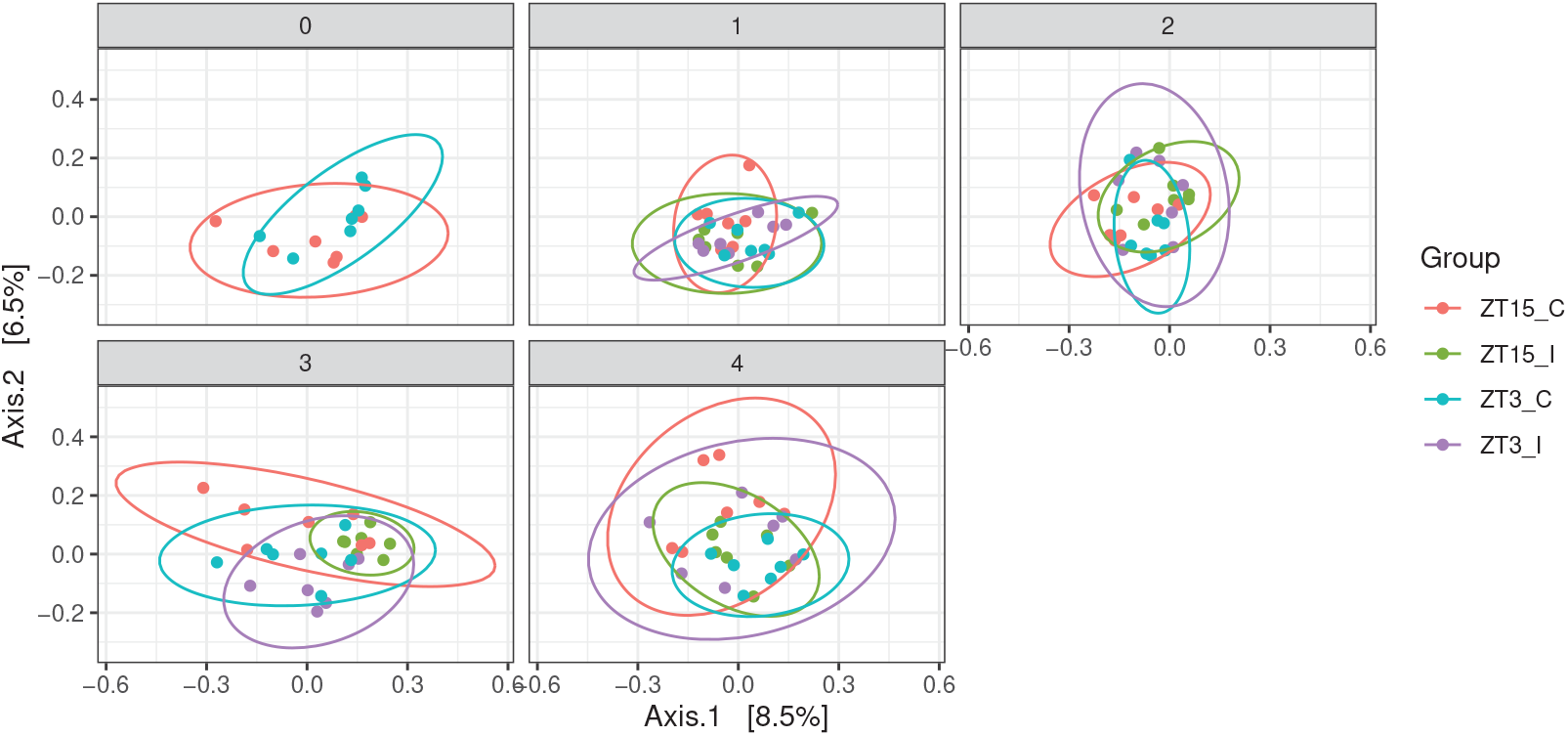
PCoA ordination plots for beta diversity metrics based on the time of infection and the period before and after infection, showing UniFrac, for vaginal samples. Each dot represents the microbial community of a sample, each color corresponds to a treatment group (ZT15_C, ZT15_I, ZT3_C, and ZT3_I). The first axis explains 8.5% of the variability and the second axis explains 6.5% of the variability in the data of vaginal samples.

We also applied PERMANOVA and betadisper to vaginal samples subset by treatment group (ZT15_C, ZT15_I, ZT3_C, and ZT3_I) to check whether the community compositions differ over time within each group (Table 2). For ZT15_C and ZT3_I samples, the independent variable Week (i.e., the week(s) before and after infection that the sample was collected) was significant for all three distance measures, but the significance with UniFrac for ZT15 C was likely due to the differences in dispersion between samples collected in week 0 and week 3 as well as samples collected one week and three weeks after infection (week 1 and week 3). For ZT15_I, the independent variable Week was significant for UniFrac and W-UniFrac but not Bray-Curtis, indicating that ZT15_I samples tend to be phylogenetically distinct among treatment groups; but the significance with UniFrac was likely due to the differences in dispersion between samples collected two and three weeks after infection as well as samples collected two and four weeks after infection. For ZT3_C, the independent variable Week was significant with Bray-Curtis and W-UniFrac but not UniFrac, suggesting that samples collected at different time points tend to have the same ASVs but difference abundance of those ASVs; the betadisper results confirmed that no significant differences in dispersion between any pair of samples in this case.

**Table 2:** PERMANOVA and betadisper results for vaginal samples subset by group. The factor here is Week. Asterisk denotes statistically significant differences (*,*p* ⩽ 0.05; **,*p* ⩽ 0.01; ***,*p* ⩽ 0.001).

| Data | Distance Metric | PERMANOVA<br>$R^2$ | $p$ | Pairs Differ in Dispersion<br>significant pair (permuted $p$) |
| --- | --- | --- | --- | --- |
| ZT15_I | Bray-Curtis | 0.068 | 0.068 | 2-3(0.040*), 2-4 (0.005**) |
|  | UniFrac | 0.070 | 0.002** |  |
|  | W-UniFrac | 0.080 | 0.046* |  |
| ZT15_C | Bray-Curtis | 0.111 | 0.002** | 0-3 (0.038*), 1-3 (0.029*) |
|  | UniFrac | 0.053 | 0.004** |  |
|  | W-UniFrac | 0.149 | 0.001*** |  |
| ZT3_I | Bray-Curtis | 0.144 | 0.003** |  |
|  | UniFrac | 0.067 | 0.002** |  |
|  | W-UniFrac | 0.103 | 0.021* |  |
| ZT3_C | Bray-Curtis | 0.084 | 0.01* |  |
|  | UniFrac | 0.023 | 0.85 |  |
|  | W-UniFrac | 0.099 | 0.012* |  |

We then subset vaginal samples by the week pre- and post-infection that the samples were collected to check whether the community compositions differ among treatment groups (Table 3). PERMANOVA and betadisper analyses confirmed that no statistically significant differences between samples of two control groups, ZT15_C and ZT3_C, collected before infection. Week 1 (one week post infection) samples were likely to have different specific ASVs but phylogenetically similar ASVs (i.e., the ASVs differ among groups are phylogenetically close to each other), since the independent variable Group was only significant with Bray-Curtis. For week 2 and week 3 samples, the independent variable Group was only significant with UniFrac, this indicated that week 2 and week 3 samples might have different rare lineages among groups. But the significance with UniFrac for week 3 samples was likely due to differences in dispersion between pairs of ZT15_I and ZT15_C, ZT15_C and ZT3_C, ZT15_C and ZT3_C. As for week 4 samples, the independent variable Group was significant with UniFrac and W-UniFrac but not Bray-Curtis, suggesting that week 4 samples tend to be phylogenetically distinct among treatment groups. Note that the significance with W-UniFrac for week 4 samples was likely due to the differences in dispersion between ZT15_I and ZT3_C samples.

**Table 3:** PERMANOVA and betadisper results vaginal samples subset by week. The factor here is Group. Asterisk denotes statistically significant differences (*,*p* ⩽ 0.05; **,*p* ⩽ 0.01; ***,*p* ⩽ 0.001).

| Data | Distance Metric | PERMANOVA | | Pairs Differ in Dispersion<br>significant pair (permuted $p$) |
| --- | --- | --- | --- | --- |
| | | $R^2$ | $p$ | |
| Week 0 | Bray-Curtis | 0.070 | 0.596 |  |
|  | UniFrac | 0.085 | 0.294 |  |
|  | W-UniFrac | 0.050 | 0.791 |  |
| Week 1 | Bray-Curtis | 0.184 | 0.001*** |  |
|  | UniFrac | 0.129 | 0.067 |  |
|  | W-UniFrac | 0.126 | 0.258 |  |
| Week 2 | Bray-Curtis | 0.081 | 0.847 |  |
|  | UniFrac | 0.183 | 0.001*** |  |
|  | W-UniFrac | 0.077 | 0.871 |  |
| Week 3 | Bray-Curtis | 0.107 | 0.449 | ZT15_I-ZT15_C (0.024*), ZT15_C-ZT3_C (0.020*)<br>ZT15_C-ZT3_I (0.028*) |
|  | UniFrac | 0.174 | 0.002** |  |
|  | W-UniFrac | 0.100 | 0.518 |  |
| Week 4 | Bray-Curtis | 0.176 | 0.062 | ZT15_I-ZT3_C (0.034*) |
|  | UniFrac | 0.146 | 0.007** |  |
|  | W-UniFrac | 0.190 | 0.037* |  |

### 3.3 Genital Tract Samples Collected Four Weeks Post-infection (Week 4 Samples)

A total of 1677 ASVs were identified across all samples collected four weeks post-infection (*n* = 92), with 107 ASVs shared among all three GTRs (occupying approximately 6.38% of all ASVs). Furthermore, those 107 shared ASVs span 6 phyla, 10 classes, and 25 orders. As shown in Fig. 6, the number of unique ASVs in Ovary/Oviduct, Uterus, Vaginal samples collected in week 4 are 531, 486, and 339, respectively.

**Figure 6:**
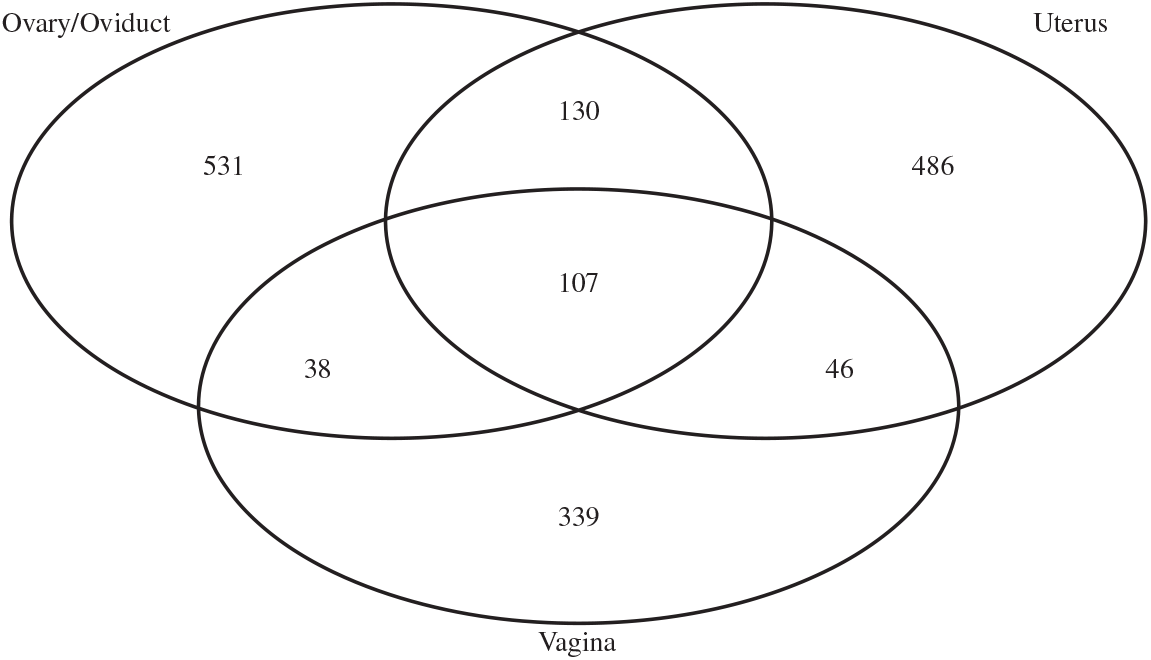
Venn diagram showing the number of shared and unique ASVs among three sites (Ovary/Oviduct, Uterus, Vagina) for the samples collected four weeks post-infection.

#### 3.3.1 Alpha-diversity of Week 4 Samples

In addition to vaginal samples, we also collected ovary/oviduct and uterus samples four weeks post-infection. The Shannon and Simpson diversity indices of week 4 samples by GTR for infection and control groups are depicted by the boxplots in Fig. C.1 in Supplementary Material. There were significant differences in both diversity indices between ovary/oviduct and vaginal samples for the control group (*p* ⩽ 0.05, Kruskal-Wallis rank-sum test), and those differences were only significant in group ZT15 C (*p* ⩽ 0.05, Kruskal-Wallis rank-sum test), as shown in Figs. 7 and C.2.

**Figure 7:**
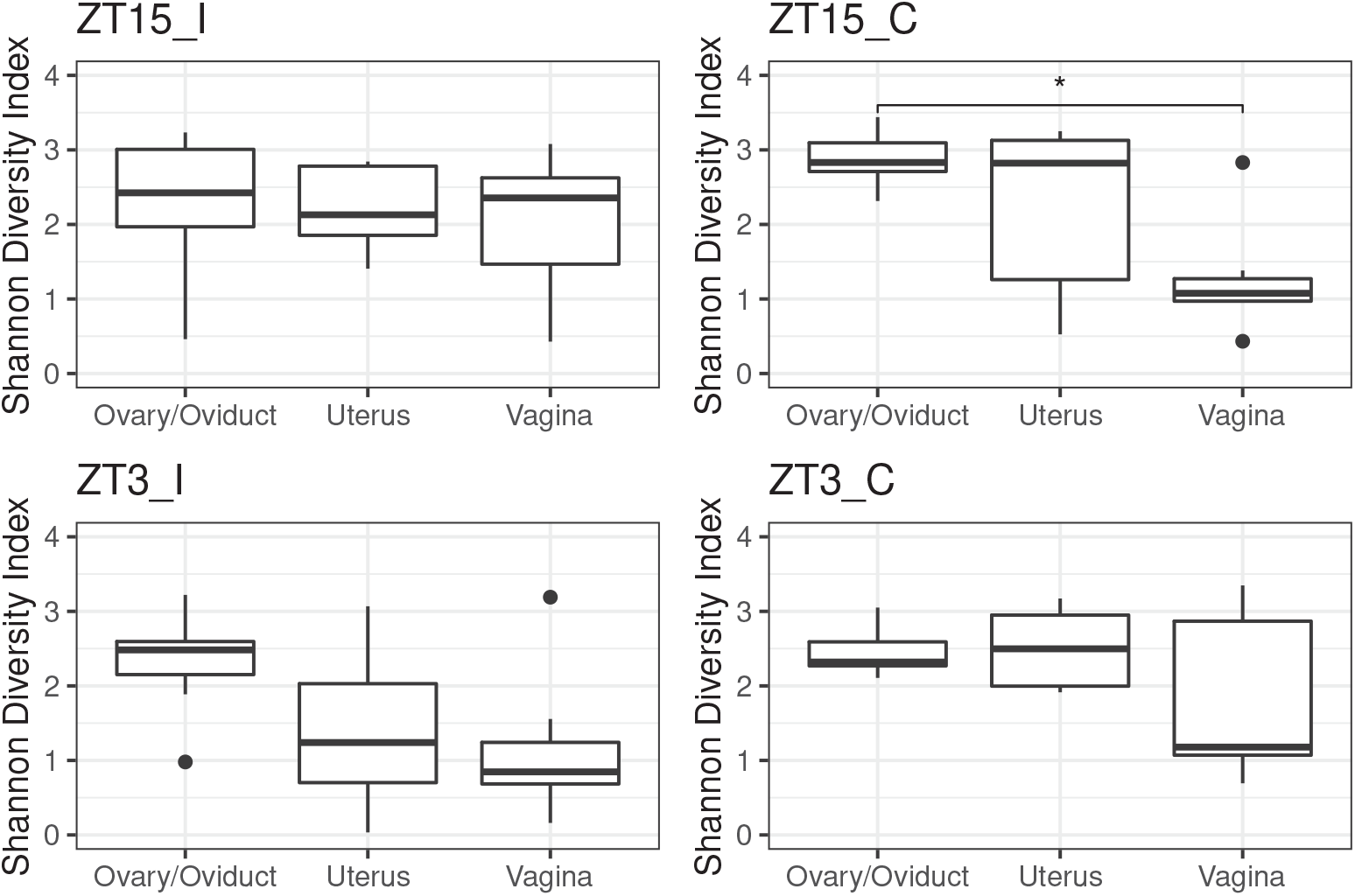
Boxplots of Shannon Index for samples collected four weeks post-infection by GTR for each group (ZT15_I, ZT15_C, ZT3_I, and ZT3_C). Statistical significance is indicated above the brackets: *,*p* ⩽ 0.05.

We also observed significant differences in Shannon diversity index between infected group (ZT15_I and ZT3_I) and control group (ZT15_C and ZT3_C) in uterin samples (*p* > 0.05, Kruskal-Wallis rank-sum test, Fig. C.3 in Supplementary Material), but the difference in uterin samples was not statistically significant (*p* > 0.05, Kruskal-Wallis rank-sum test) when comparing among individual groups (ZT15_I, ZT3_I, ZT15_C and ZT3_C), as shown in Fig. 8.

**Figure 8:**
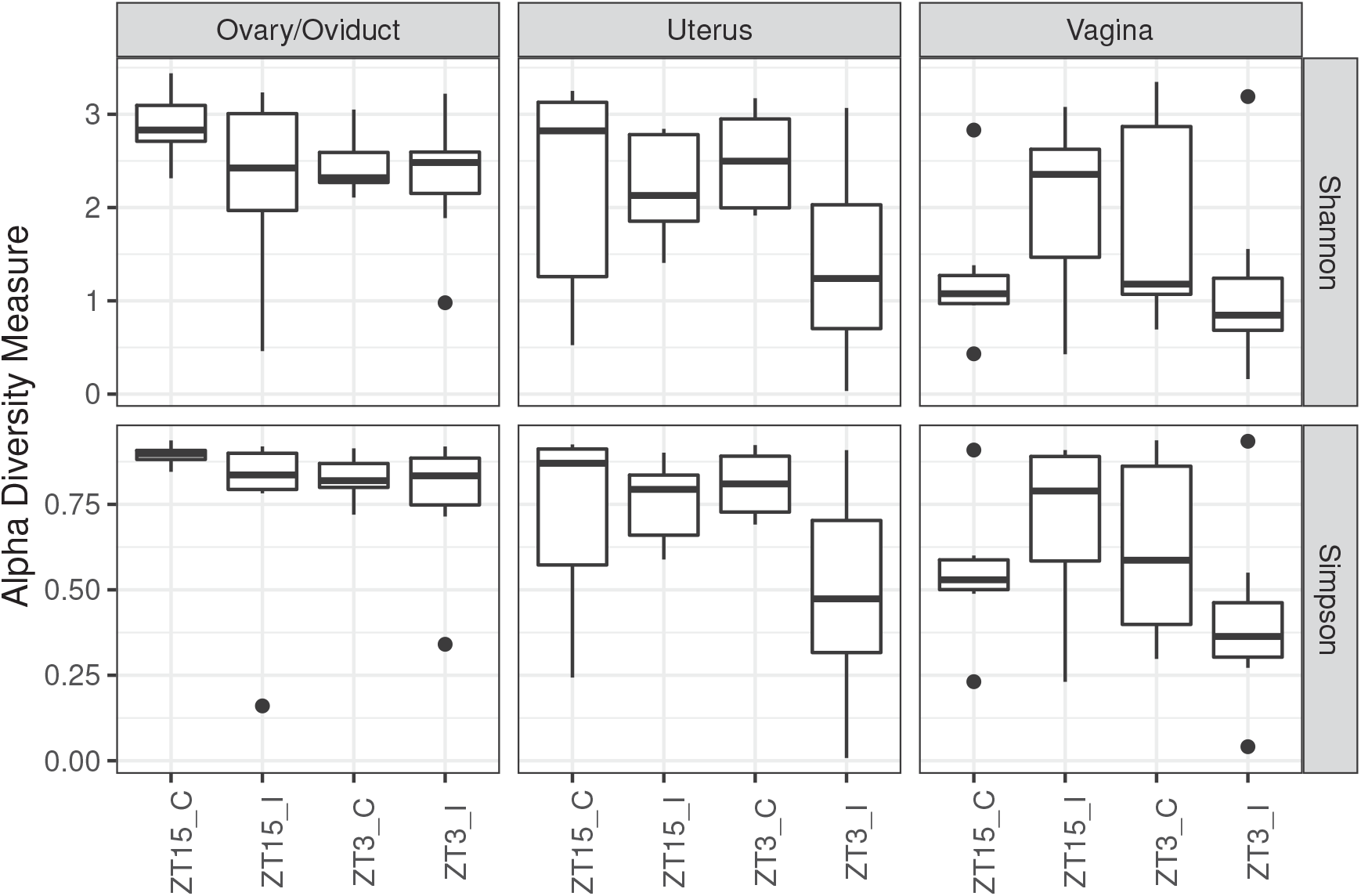
Boxplots of Shannon and Simpson Index for samples collected four weeks post-infection by group (ZT15_I, ZT15_C, ZT3_I, and ZT3_C) per GTR.

#### 3.3.2 Beta Diversity of Week 4 Samples

For all samples collected four weeks post-infection (week 4), PERMANOVA and betadisper confirmed that both the independent variables GTR and Group had significant impact on the microbiota composition with all three distance measures we tested (Table 4). The PCoA ordination plots for week 4 data faceting by group and GTR are shown in Figs. D.1 and D.2 in Supplementary Material, respectively. We also performed CCA including both independent variables Group and GTR, and confirmed that both independent variables were significant (*p* = 0.001) and the first three constrained axes were significant (*p* = 0.001 for first two constrained axes, *p* = 0.002 for the third constrained axis). The CCA ordination diagrams show that samples from some groups were tightly cluster together within certain GTR (Supplementary Fig. E.1).

**Table 4:** PERMANOVA results for all samples collected four weeks post-infection (week 4). Asterisk denotes statistically significant difference (*,*p* ⩽ 0.05; **,*p* ⩽ 0.01; ***,*p* ⩽ 0.001). Note that betadisper did not identify any pair with statistically significant differences in dispersion.

| Distance Metric | Factor | PERMANOVA |  |
| --- | --- | --- | --- |
| | | $R^2$ | $p$ |
| Bray-Curtis | GTR | 0.138 | 0.001*** |
|  | Group | 0.089 | 0.001*** |
| | GTR × Group | 0.087 | 0.003** |
| Unifrac | GTR | 0.070 | 0.001*** |
|  | Group | 0.048 | 0.001*** |
| | GTR × Group | 0.071 | 0.032* |
| W-UniFrac | GTR | 0.163 | 0.001*** |
|  | Group | 0.072 | 0.004** |
| | GTR × Group | 0.100 | 0.005** |

We then subset week 4 samples by group to examine the effect of GTR on community compositions (Table 5). For ZT15_I samples, the independent variable GTR is significant for Bray-Curtis and Unifrac but not W-Unifrac, which suggests that ZT15_I samples have different ASVs based on the genital tract region and likely to be phylogenetically distant in less finer niches. For samples in other three groups (ZT15_C, ZT3_I, and ZT3_C), the independent variable GTR is significant for all three distance metrics tested; but for ZT15_ with Bray-Curtis and W-UniFrac, the differences were likely due to differences in dispersion between ovary/oviduct and vagina. CCA carried out on samples collected from each genital tract region confirmed that both the constrained ordination and the independent variable Group were significant for samples within each genital tract region (*p* = 0.001 for ovary/oviduct and uterus samples and *p* = 0.014 for vaginal samples). The significance level of individual axis is different among GTRs: for ovary/oviduct samples, the first two constrained axes were significant (*p* = 0.001 and *p* = 0.014 for the first and second constrained axis, respectively); for uterus samples, the first constrained axis was significant (*p* = 0.001); for vaginal samples, the first constrained axis was marginally significant (*p* = 0.066). Fig. 9 shows the CCA ordination diagram with first two constrained axes for samples within each genital tract region. We can see a clear separation among different treatment groups in ovary/oviduct samples in both constrained axes, we can also see a clear separation between ZT15_C and other three groups in uterus samples along the first constrained axis. In addition, we summarized how ASVs within each phylum was represented along those significant constrained axes for ovary/oviduct (Fig. 10) and uterus samples (Fig. 11), respectively. The ASVs that cluster unusually from the rest of their phylum along the constrained axis are labeled to the family level. Take ovary/oviduct samples as an example (Fig. 10), some ASVs in the *Deinococcaceae* family within the *Deinococcota* phylum were positioned mostly in the positive CCA1 direction toward the ZT15_I samples; some ASVs in the *Myxococcaceae* family within the *Myxococcota* phylum were positioned mostly in the negative CCA2 direction toward the ZT15 C samples; some ASVs in the *Obscuribacteraceae* family within the *Cyanobacteria* were positioned mostly in the positive CCA2 direction toward the ZT3_I samples.

**Table 5:**
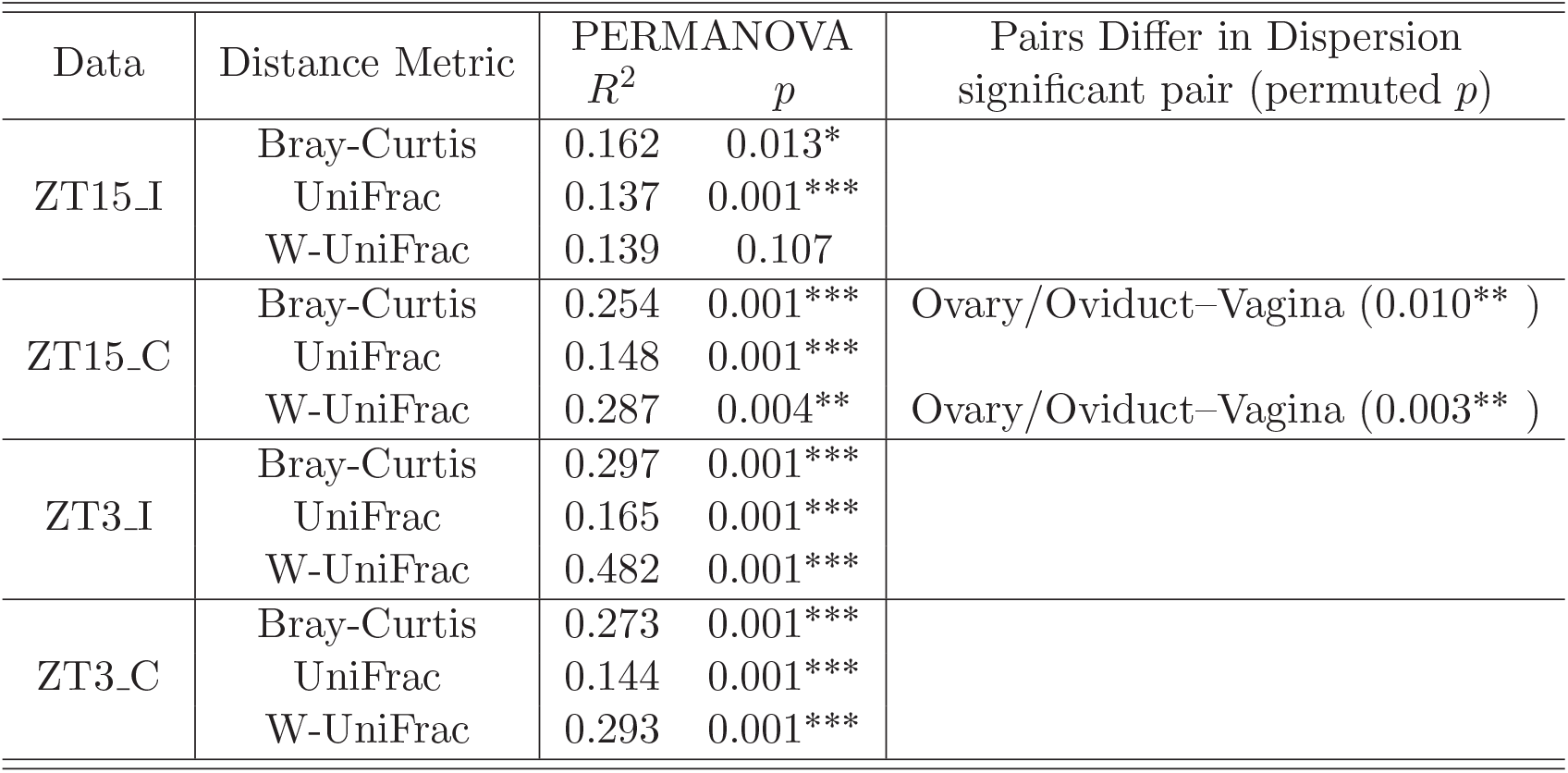
PERMANOVA and betadisper results for samples collected four weeks post-infection (week 4). The independent variable here is GTR. Asterisk denotes statistically significant difference (*,*p* ⩽ 0.05; **,*p* ⩽ 0.01; ***,*p* ⩽ 0.001).

**Figure 9:**
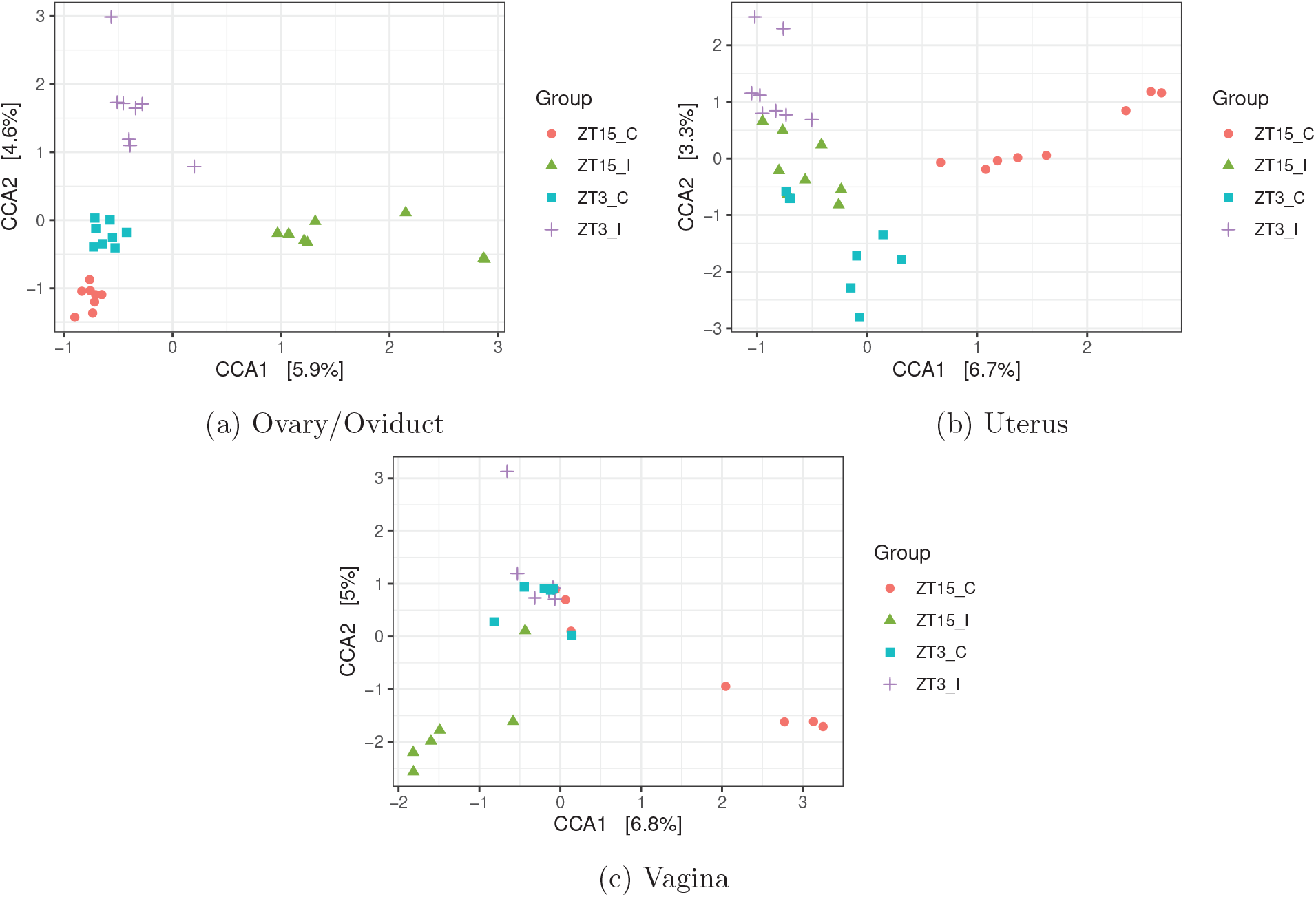
Ordination diagram with first two constrained axes for (a) Ovary/Oviduct samples, (b) Uterus samples, and (c) Vaginal samples collected four weeks post-infection (week 4). The first and second constrained axis explains 5.9% and 4.6% of the constrained inertia in the Ovary/Oviduct samples, respectively. For Uterus samples, the first and second constrained axis explains 6.7% and 3.3% of the constrained inertia in the data, respectively. For Vaginal samples collected four weeks post-infection (week 4), the first and second axis explains 6.8% and 5% of the constrained inertia in the data, respectively.

**Figure 10:**
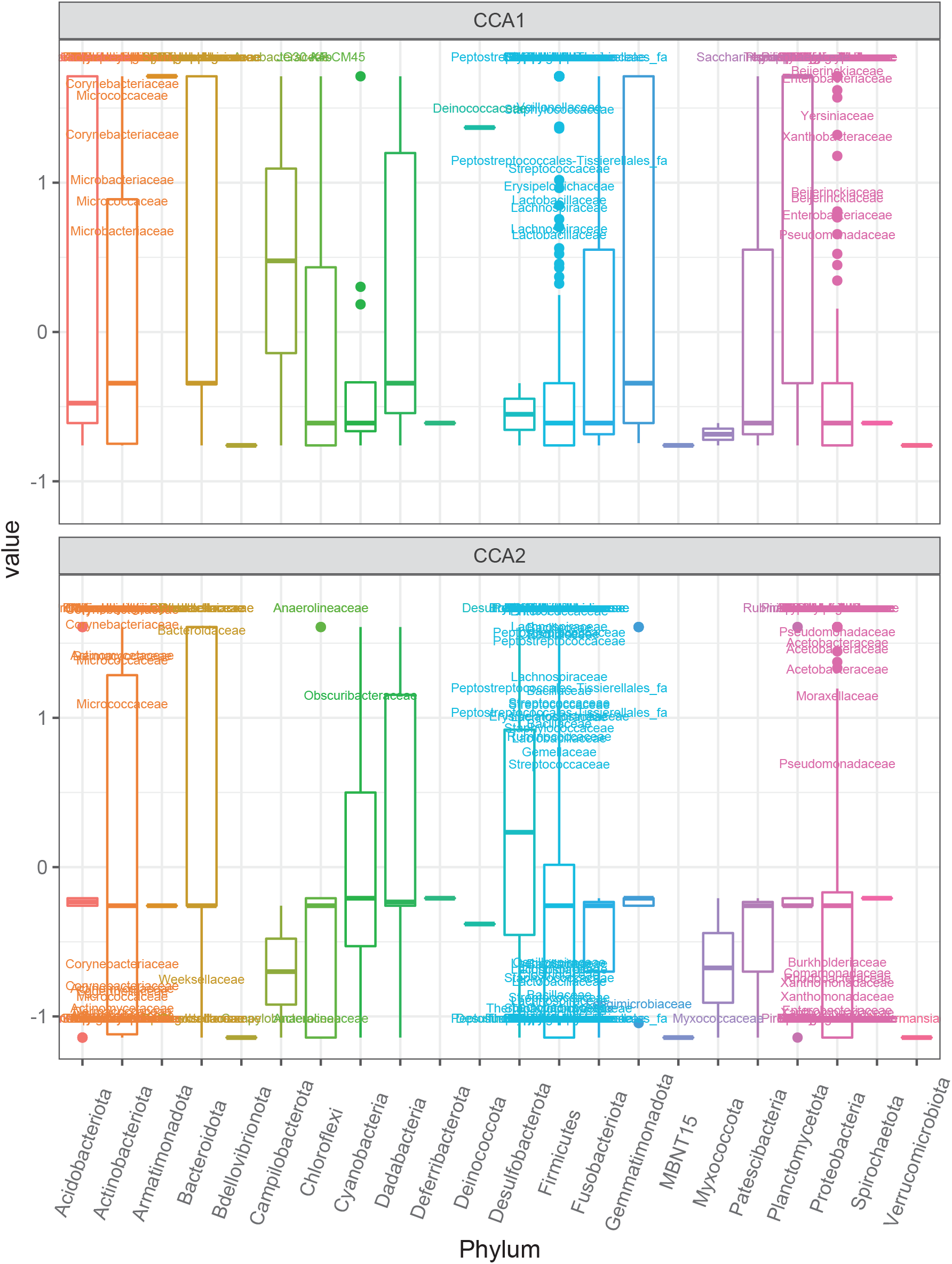
Boxplot of ASVs of the first two constrained axes from the CCA with Ovary/Oviduct samples collected four weeks post-infection (week 4), shaded/separated by phylum. Within each phylum, only the ASVs with CCA1>0.5 or CCA2>0.5 or CCA2<-0.75 are labeled to the family level.

**Figure 11:**
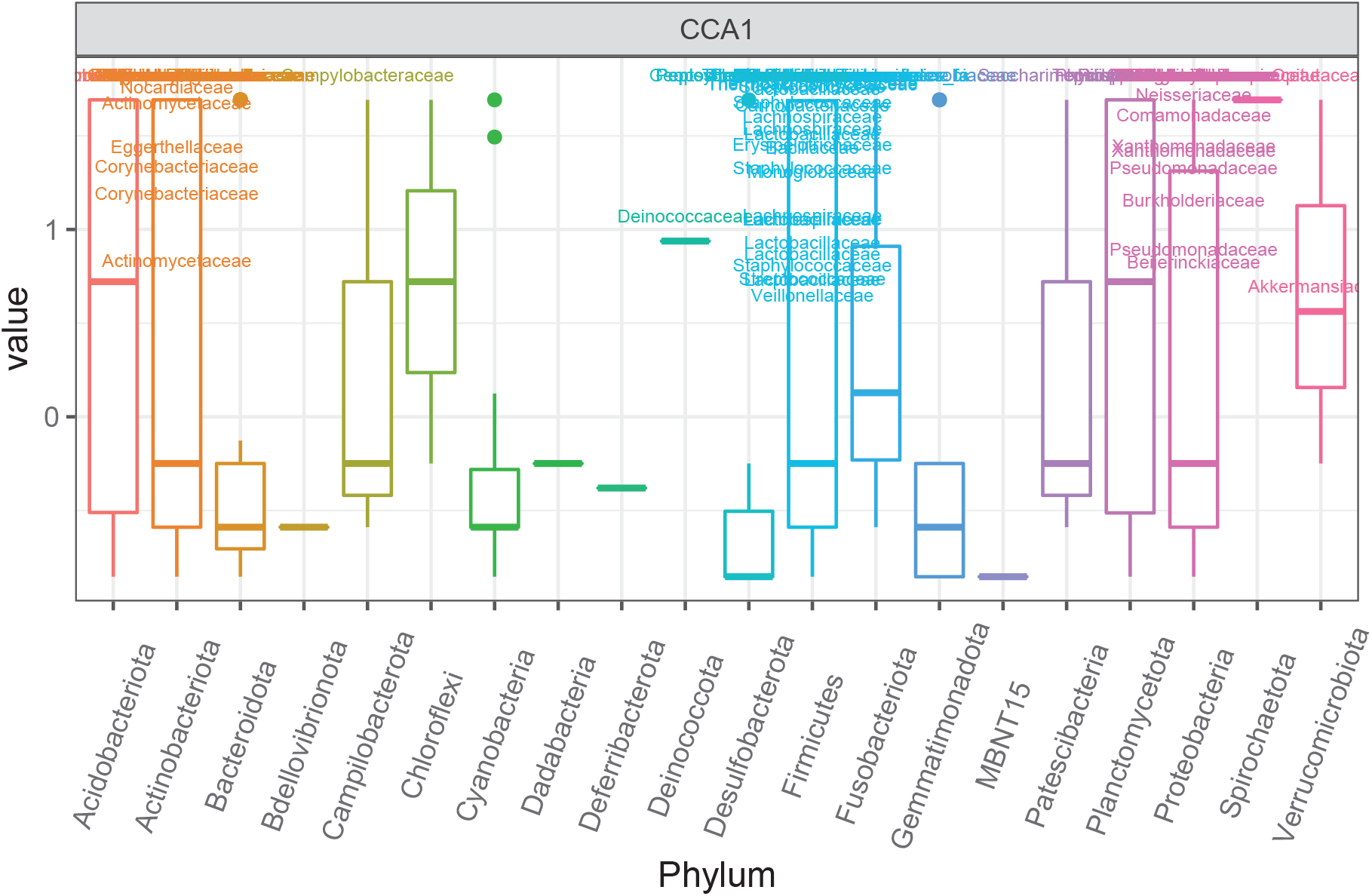
Boxplot of ASVs of the first constrained axis from the CCA with Uterus samples collected four weeks post-infection (week 4), shaded/separated by phylum. Within each phylum, only the ASVs with CCA1>0.5 are labeled to the family level.

We also checked whether the community composition differ among groups within each GTR (Table 6). The independent variable group was significant with all three distance measures for ovary/oviduct and uterus samples; but the differences for ovary/oviduct samples might due to the differences in dispersion between ZT15_I and ZT15_C (Bray-Curtis) or ZT15_I and ZT3_I (UniFrac). As for vaginal samples, the independent variable Group was significant for both Unifrac and W-Unifrac but not Bray-Curtis, this indicated that vainal microbial communities tend to be phylogenetically distinct among treatment groups. The significant differences in dispersion between ZT15_I and ZT3_C with UniFrac might be the cause of the differences among groups in vaginal samples collected four weeks post-infection (week 4). We also used CCA to assess the association between microbiota composition and GTR within each group (ZT15_I, ZT15_C, ZT3_I, ZT3_C). Both the constrained ordination and the independent variable GTR were confirmed to be significant within each group (*p* = 0.008 for ZT15_I and *p* = 0.001 for the other three groups). And for all groups, only the first constrained axis was significant (*p* = 0.005 for ZT15_I and *p* = 0.001 for the other three groups). From CCA ordination diagrams (Fig. 12) we can see that the first constrained axis was mostly separating vaginal samples from ovary/oviduct samples, and the separation was clear without any overlap for ZT15_I, ZT15_C, and ZT3_C. We also summarized how ASVs in each phylum were represented along the first constrained axis within each group in Fig. 13, with candidate indicator ASVs (positioned toward vaginal samples) being labeled to the family level. For example, the phylum *Chloroflexi* was present in both ZT15_I and ZT3_C samples but not in ZT15_C or ZT3_I samples; within this phylum, some ASVs in the family *Anaerolineaceae* were positioned mostly in the positive CCA1 direction towards vaginal samples in group ZT15_I. For the phylum *Actinobacteriota*, some ASVs in the family *Dermacoccaceae* were positioned mostly towards the positive CCA1 direction in vaginal samples of groups ZT15_I, ZT15_C, and ZT3_C but not ZT3_I.

**Table 6:**
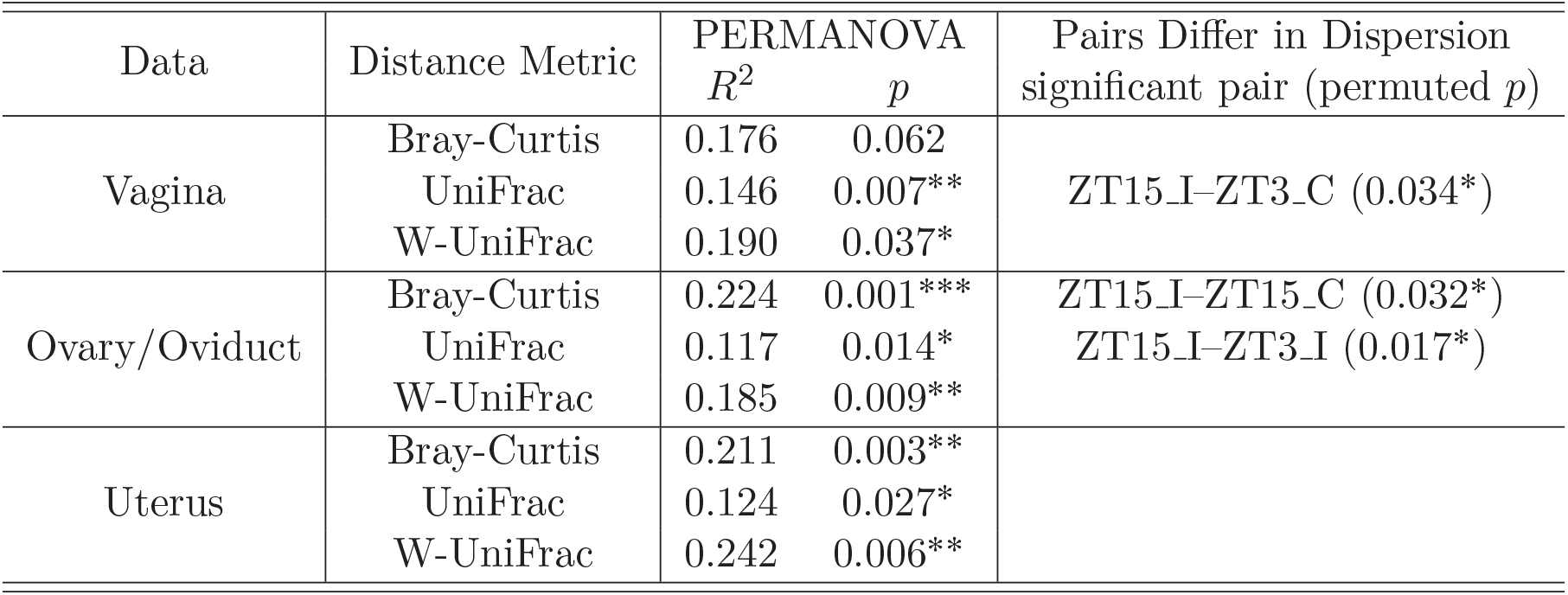
PERMANOVA and betadisper results for samples collected four weeks post-infection (week 4). The independent variable here is Group. Asterisk denotes statistically significant difference (*,*p* ⩽ 0.05; **,*p* ⩽ 0.01; ***,*p* ⩽ 0.001).

| Data | Distance Metric | PERMANOVA<br>$R^2$ | $p$ | Pairs Differ in Dispersion<br>significant pair (permuted $p$) |
| --- | --- | --- | --- | --- |
| Vagina | Bray-Curtis | 0.176 | 0.062 |  |
|  | UniFrac | 0.146 | 0.007** | ZT15_I-ZT3_C (0.034*) |
|  | W-UniFrac | 0.190 | 0.037* |  |
| Ovary/Oviduct | Bray-Curtis | 0.224 | 0.001*** | ZT15_I-ZT15_C (0.032*) |
|  | UniFrac | 0.117 | 0.014* | ZT15_I-ZT3_I (0.017*) |
|  | W-UniFrac | 0.185 | 0.009** |  |
| Uterus | Bray-Curtis | 0.211 | 0.003** |  |
|  | UniFrac | 0.124 | 0.027* |  |
|  | W-UniFrac | 0.242 | 0.006** |  |

**Figure 12:**
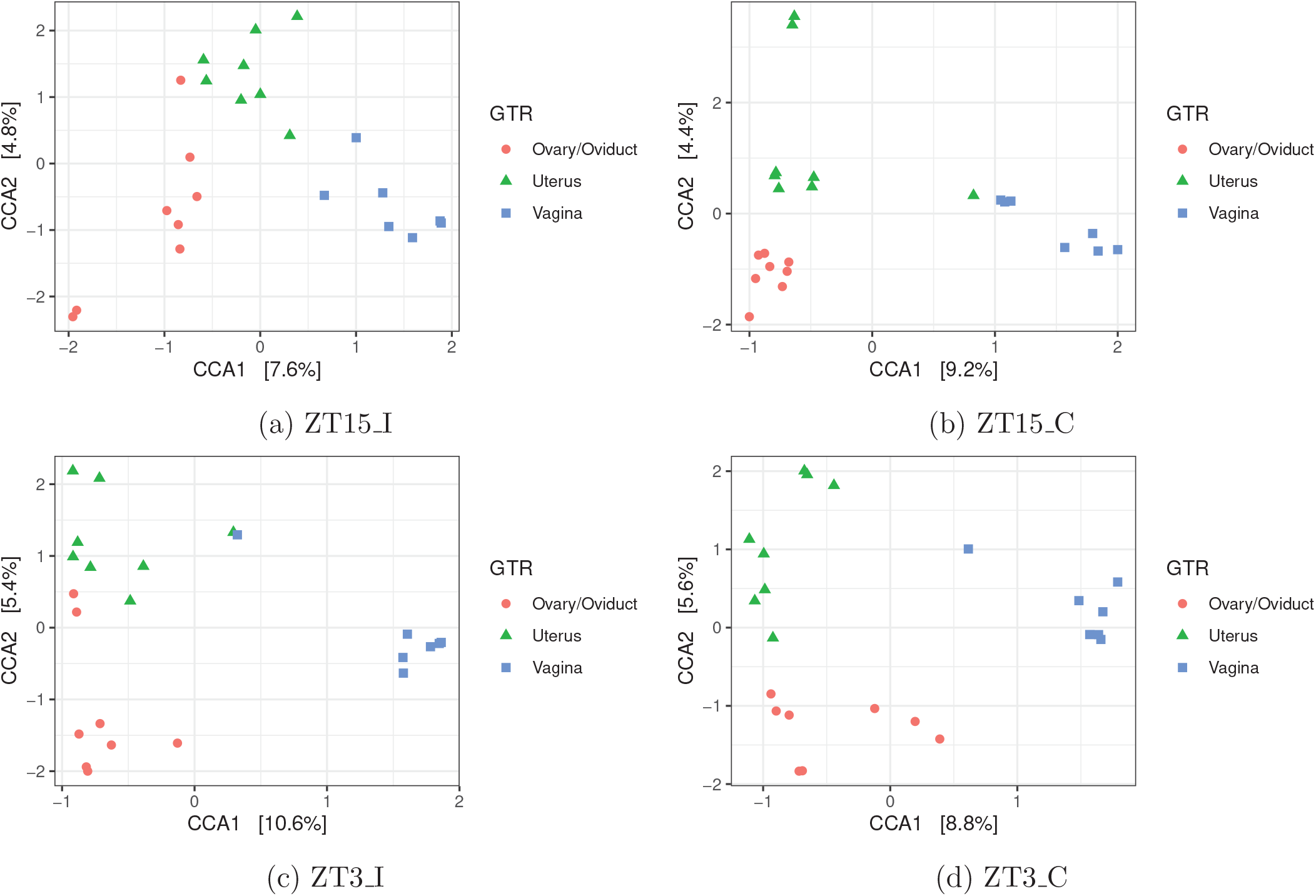
Ordination diagram with first two constrained axes for (a) ZT15_I, (b) ZT15_C, (c) ZT3_I, and (d) ZT3_C group samples collected four weeks post-infection (week 4). The first constrained axis explains 7.6%, 9.2%, 10.6% and 8.8% of the constrained inertia in the data of group ZT15_I, ZT15_C, ZT3_I, and ZT3_C, respectively.

**Figure 13:**
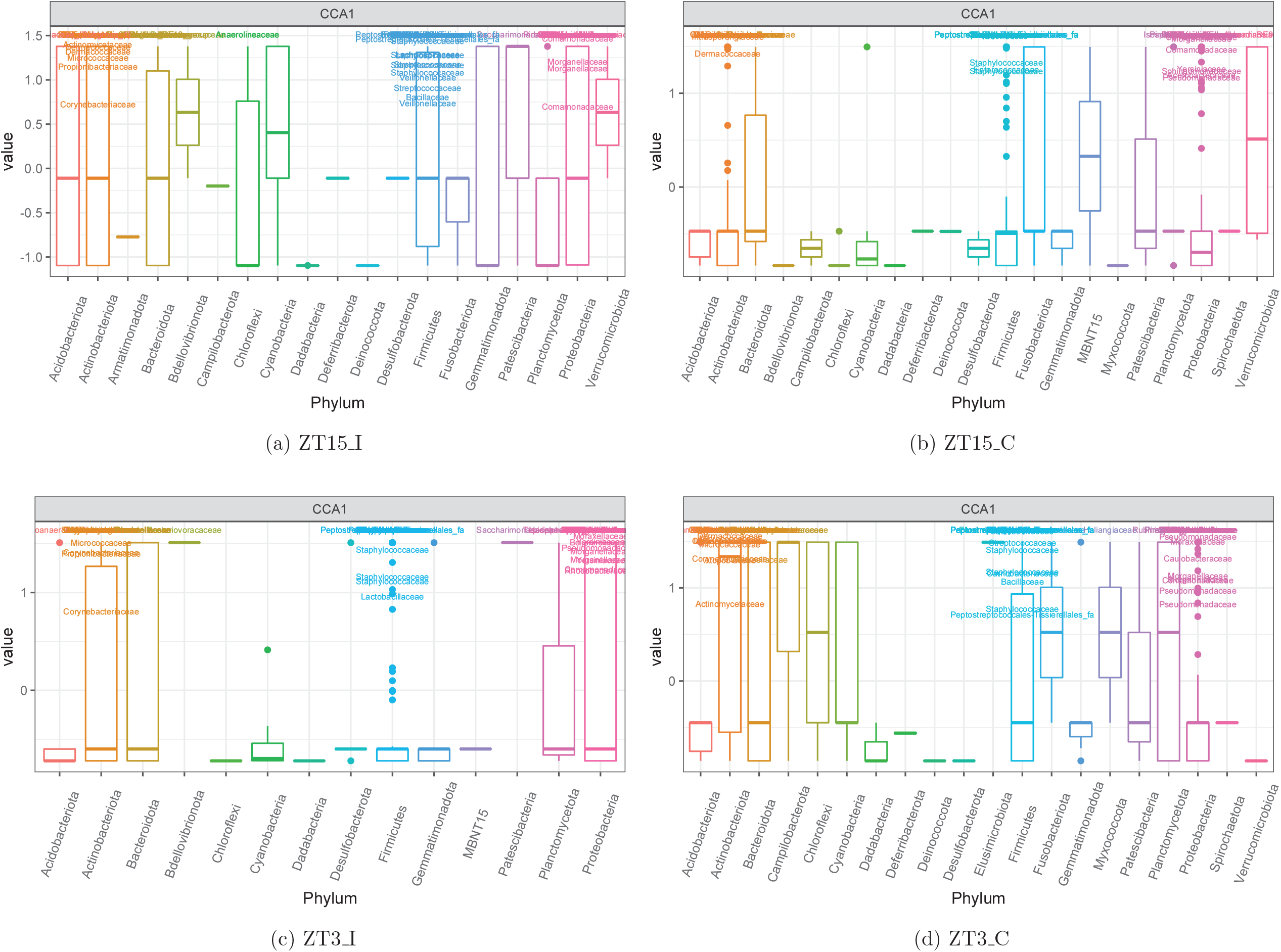
Boxplot of ASVs of the first constrained axis from the CCA with (a) ZT15_I, (b) ZT15 C, (c) ZT3_I, and (d) ZT3_C samples collected four weeks post-infection (week 4), shaded/separated by phylum. Within each phylum, only the ASVs with CCA1>0.5 are labeled to the family level in (a), (c) and (d); only the ASVs with CCA1> 1 are labeled in (b). Note that in (a), (b) and (d) the ASVs labeled here can separate vaginal samples from others; while in (c) the ASVs labeled here can separate 6 out of 7 vaginal samples from the others.

A total of 23 phyla were observed across all samples collected four weeks post-infection with most ASVs classfied to one of the four phyla: *Actinobacteriota, Bacteroidota, Firmicutes*, and *Proteobacteria*. We observed variations among individuals within each group and the genital tract region, but for in the group ZT3_I, *Firmicutes* was the dominant phyla in uterus samples (Fig. 14).

**Figure 14:**
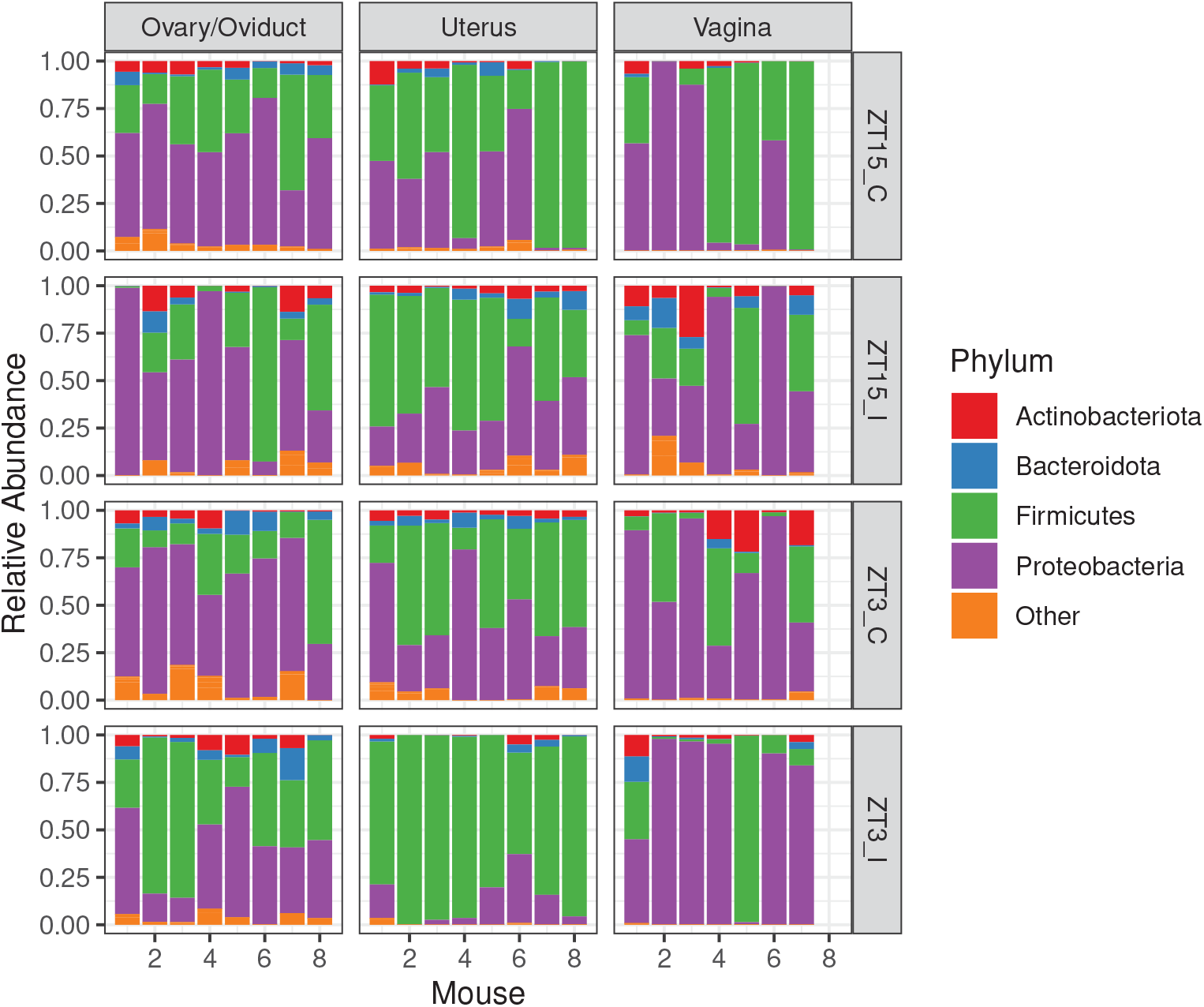
Phylum-level comparison of relative abundance of ASVs in samples collected four weeks post-infection, faceted by group and GTR. All phyla with median relative abundance across all samples ⩽0.01 are grouped as “Other”.

## 4 Discussion

We have previously shown that the time of infection was crucial in determining *Chlamydial* pathogenicity [14]. Mice infected with *Chlamydia* at ZT3, early rest period, had more infectivity and pathology than mice infected at ZT15, early active period. We still do not understand the mechanism underlying these differences in the pathogenic outcome of *Chlamydia* infection. To determine what processes might be involved in this interesting phenomenon, we investigated the genital tract microbiome’s role from the vagina to the uterus and ovary/oviduct of *Chlamydia*-infected mice.

The microbiome association with *Chlamydia* infection has been extensively studied, mainly in women [30, 31, 17, 24, 32]. Mice, guinea pigs, and other animal models might not be the best for studying the human vaginal microbiome [33] since the vagina is alkaline in mice and acidic for humans [34, 35, 36]. However, we have expanded the sites or parts of the mice genital tract to examine the microbiome in this study. The mice model has been extensively used in determining *Chlamydial* pathogenesis by most investigators in the field [37, 38, 14]. In addition, we reported the effect of infection at different times of the day using a mice model [14]. That is why we went ahead and used the mice model to understand the possible role of time of infection on the genital tract microbiome. Most *Chlamydia* associated studies in women analyze the vaginal microbiome. We extended from the vagina through the uterus to the oviduct/ovary. This study highlights that the time of infection is essential in determining the richness and diversity of microbiome present in the vagina. The richness and diversity varied with the time of infection; however, this change was not significant. Microbiome collected in the third and fourth-week post-infection had the most variation. However, the sham-infected control mice had more variation over time than the *Chlamydia* infected mice. There was a significant difference in the richness and diversity of the vaginal microbiome with the time of infection. Overall, mice not infected with *Chlamydia* had the most variation in the vaginal microbiome, implying that the microbiome was highly variable in the natural process without competition from *Chlamydia*. When we parsed the infection into the infection time, at four weeks post infection, mice infected at ZT15 showed more variation than mice infected at ZT3. The sham-infected control mice at ZT3 had more richness and variation than the *Chlamydia* infected mice. In contrast, infection at ZT15 increased richness and variation, showing that the time of infection had different microbiome outcomes. This result was verified using PCoA analysis, which showed that the vaginal microbial community composition was different for infection times, and this varied with the duration of infection. Differential tree (Fig. B.1 in Supplementary Material) portrayed the diverse microbial community present in the vagina during the course of infection.

We also examined the microbiome of other regions of the genital tract: the uterus and the ovary/oviduct four weeks post-infection. This experiment was based on the premise that the microbes are found in most parts of mammals, including regions usually thought to be immunocompetent and protected from microbes [39, 40, 41]. It has been reported that the placenta and other parts of the human reproductive tract have their microbiome [42, 43, 44]. Our data showed that the uterus and ovary/oviduct’s microbiome had bacteria unique from the vagina and shared some bacteria with the vagina (Fig. 6). This result is interesting as most of the studies have solely focused on studying the vaginal microbiome [45, 46, 47, 17]. Understanding the role of the microbial community in the upper genital tract during *Chlamydia* infection is necessary since *Chlamydia* must climb up the genital tract to cause its pathology in the upper genital tract.

The time of *Chlamydia* and sham infection was crucial in determining the richness and diversity of the ovary/oviduct, uterus, and vaginal microbiome during *Chlamydia* infection. The microbiome variation between different regions of the genital tract in mice infected with *Chlamydia* at ZT15 was less than mice infected at ZT3. On the other hand, the control mice had significant variation at ZT15. When we compared the different regions of the genital tract by infection time, only the uterus samples showed a considerable difference in the microbiome between the two treatment groups. This result signifies that focusing on just the vaginal microbiome in understanding what is going on in the female reproductive tract during *Chlamydia* infection is not enough. The vaginal microbiome tells us what happens at the point of infection and not necessarily in the upper genital tract. The vaginal microbiome might not be associated with upper genital tract pathology. These microbial communities of the different regions of the genital tract are diverse, and these differences are accentuated based on the time of infection. This was visualized via PCoA analysis. The ovary/oviduct and uterus microbial communities were distinctly demarcated compared to the vaginal microbial communities when comparing the time of infection, showing that infection time was critical in delineating the microbes present in these regions compared to the vagina. We considered that the microbiome present in other areas in the genital tract might be more critical in predicting *Chlamydial* pathogenicity than just looking at the vaginal microbiome.

Based on the context that the microbes in the ovary/oviduct were demarcated based on infection time, we looked at the species present in that region. We noticed that *Actinobacteria, Cyanobacteria, Firmicutes*, and *Proteobacteria* were the dominant phyla. In addition, we observed that different phyla were present in the uterus compared to the ovary/oviduct samples. This result was expected based on the earlier result that the characteristics of the ovary/oviduct were different based on the time of infection. When we put all the bacteria from all regions of the genital tract together, we showed that the bacteria were not the same based on the time of infection. This might be due to the influence of the microbiome in the ovary/oviduct. *Proteobacteria, Firmicutes*, and *Actinobacteria* were the top three phyla found in these regions. *Firmicutes* and *Proteobacteria* are predominant phyla in both the mouse and human gut microbiome [48, 49, 50]. The ratio of *Firmicutes* with other bacteria has been associated with the pathogenesis of irritable bowel syndrome and obesity [49, 51, 52].

## 5 Conclusions

Overall, this study showed that the time of infection is essential in determining the bacterial population present in the microbiome of the different regions of the genital tract. We do not fully understand what is responsible for this phenomena and how the time of infection is essential in determining which ASVs are present in different regions of the genital tract. However, this difference appears to be more accentuated in the ovary/oviduct and uterus microbiome than in the vaginal microbiome. Most studies have focused on the vaginal microbiome and not the microbiome found in other parts of the reproductive tract. We show that it might be essential to analyze the microbiome in the different parts of the genital tract to provide a complete picture. Also, we are proposing that the mice model can determine the effect of *Chlamydia* infection on the microbiome. This assertion is because of the similarity we observed between the upper genital tract microbiome and the gut microbiome. The gut microbiome has been associated with disease pathogenesis [49, 51, 52]. Note that the upper genital tract undergoes deleterious changes during *Chlamydia* infection. In this study, we focused on the time of infection since we have previously shown the infection and pathogenesis were different based on the time of infection.

## Acknowledgements

We thank Ming Tan, German Enciso, and Arthur Lander for discussions and feedback on the research topic as well as Audrey Fu and Ellie Mokhtari for helpful discussions on data analysis. We would also like to thank the technicians at Morehouse School of Medicine, Center for Laboratory Animal Resources for their prompt and attentive service.

## Availability of data and materials

The data that support the findings of this study are available from the corresponding author, LZ, upon reasonable request.

## Ethics approval and consent to participate

### Animal Protocol Approval Statement

This study was carried out in strict adherence to the Guide for the Care and Use of Laboratory Animals of the National Institutes of Health recommendations. The Institutional Animal Care and Use Committee (IACUC) of Morehouse School of Medicine approved the study protocol (Protocol Number: 16–24).

## Consent for publication

Not applicable

## Authors’ contributions

LZ and YOO conceived, designed, and supervised the experiments; SRL and YOO performed the experiments; SRL and YOO performed the animal work; SRL and YOO assisted with data analysis on *Chlamydia* infectivity; LZ conducted the bioinformatic analyses and statistical analyses; LZ, SRL, and YOO wrote the original draft of the manuscript; SRL, FOE, JUI, LZ and YOO contributed to the manuscript writing, editing, and revision.

