## Supplemental Figures for "Genital tract microbiome dynamics are associated with time of *Chlamydia* infection"

#### A Alpha Diversity for Vaginal Samples

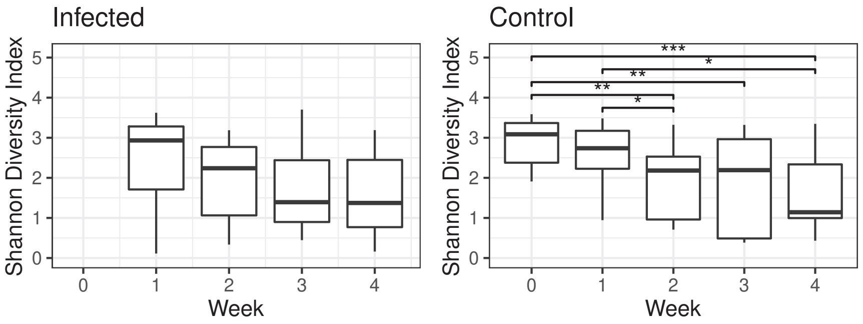


(a) Shannon Diversity Index

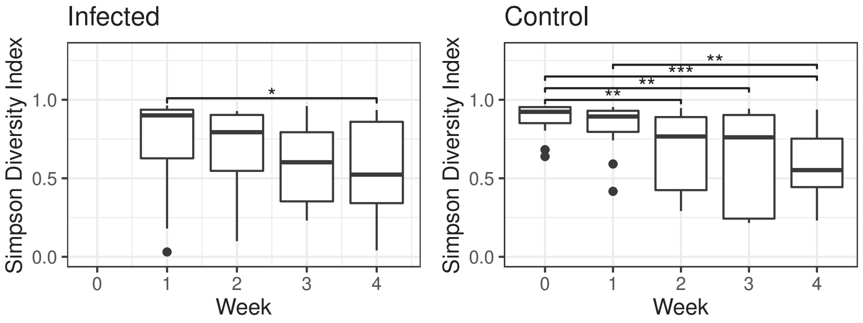


(b) Simpson Diversity Index

Figure A.1: Boxplots of Shannon (panel (a)) and Simpson Diversity Index (panel (b)) of vaginal samples in infected group (ZT15_I and ZT3_I) and control group (ZT15_C and ZT3_C) over time. The upper and lower whiskers extend from the hinge to the largest or smallest value no further than 1.5×IQR (inter-quartile range, the distance between the first and third quartiles), respectively. Statistical significance is indicated above the brackets: $*, p \leqslant 0.05$; $**, p \leqslant 0.01$; $***, p \leqslant 0.001$ (Wilcoxon rank sum test).

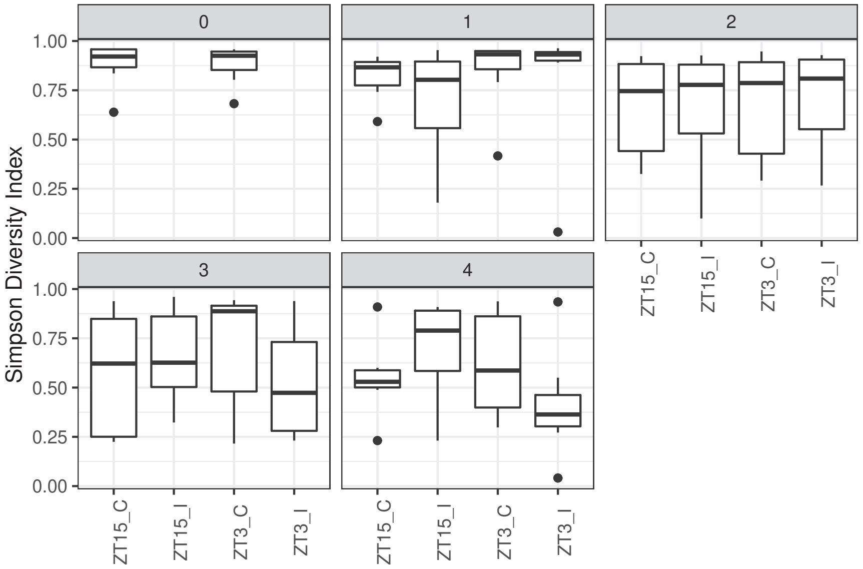


Figure A.2: Boxplots of Simpson Diversity Index for vaginal samples by group (ZT15_C, ZT15_I, ZT3_C, and ZT3_I) per week.

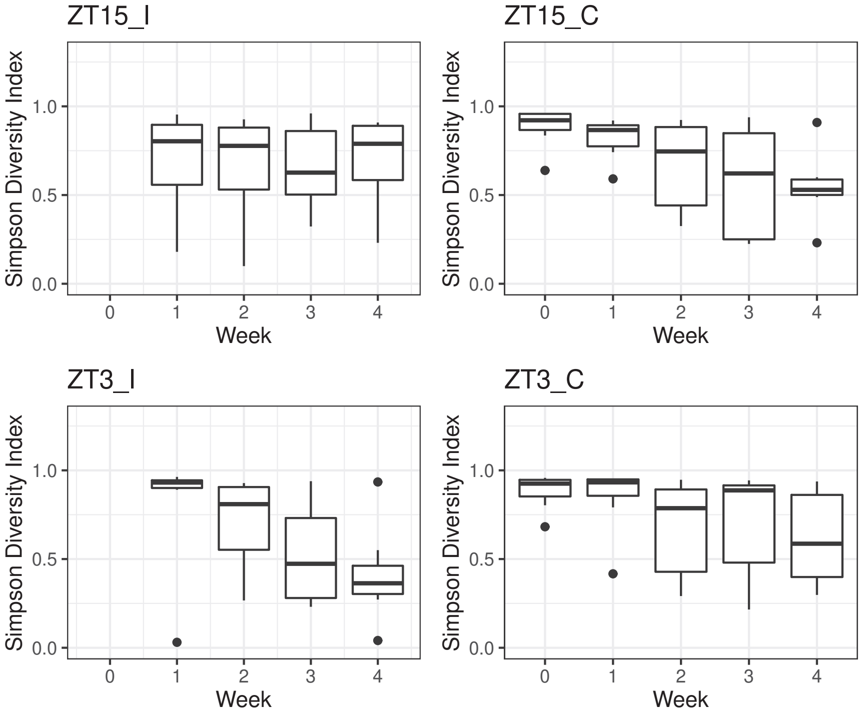


Figure A.3: Boxplots of Simpson Diversity Index for vaginal samples per group (ZT15_C, ZT15_I, ZT3_C, and ZT3_I) over time.

### B Differential Heat Tree for Vaginal Samples

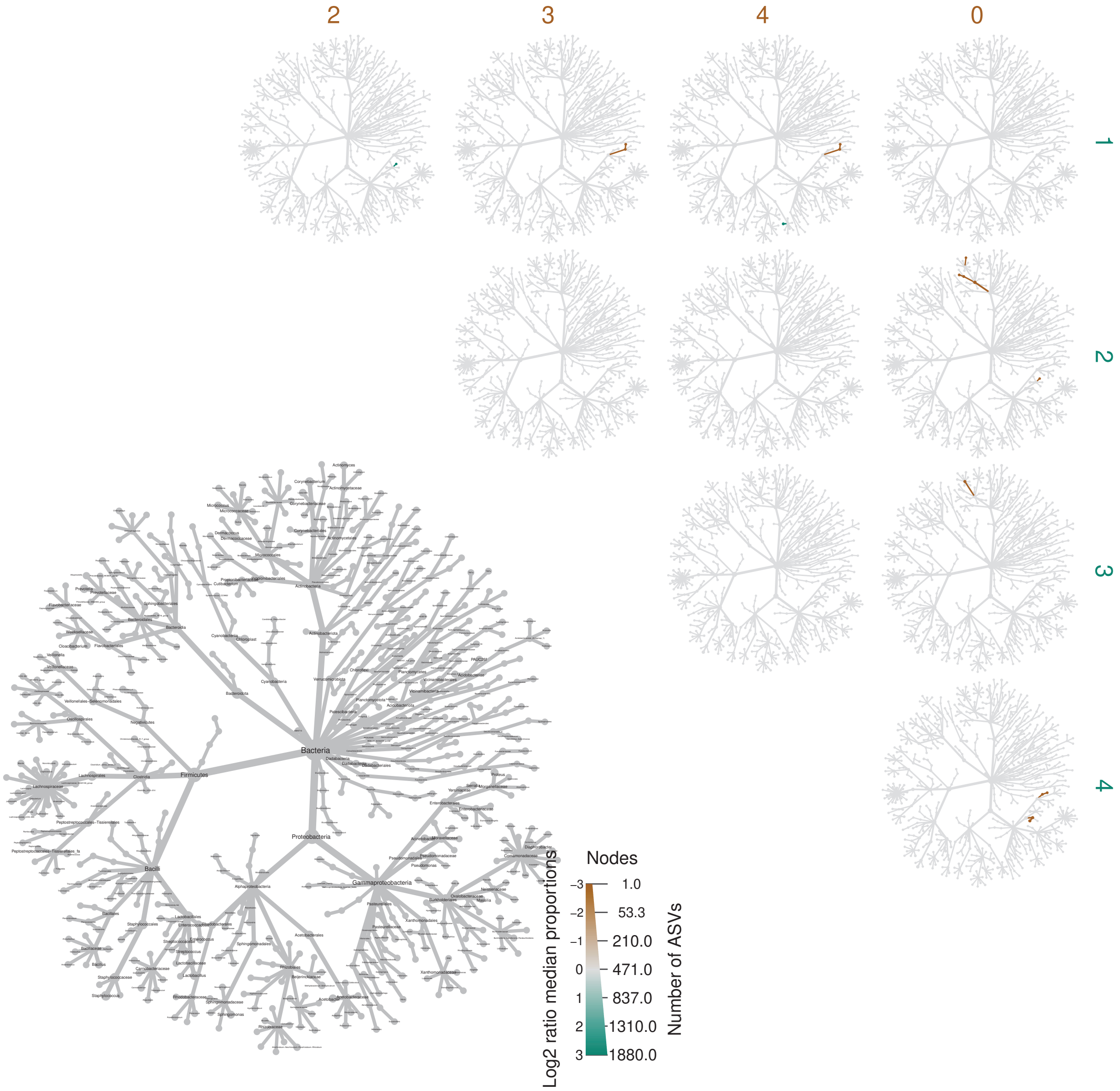


Figure B.1: Differential heat trees based on pairwise comparison of ASV relative abundances of vaginal samples collected before and post infection. The bottom left taxonomic tree acts as a reference and represents all genera present in vaginal samples. All colored taxa are significantly different between sampling weeks (Wilcoxon Rank Sum test is used with FDR correction $p < 0.05$). Color intensity corresponds to the log of ratio of median abundances in those two groups being compared, and node size corresponds to the number of ASVs of each taxon.

### C Alpha Diversity for Week 4 Samples

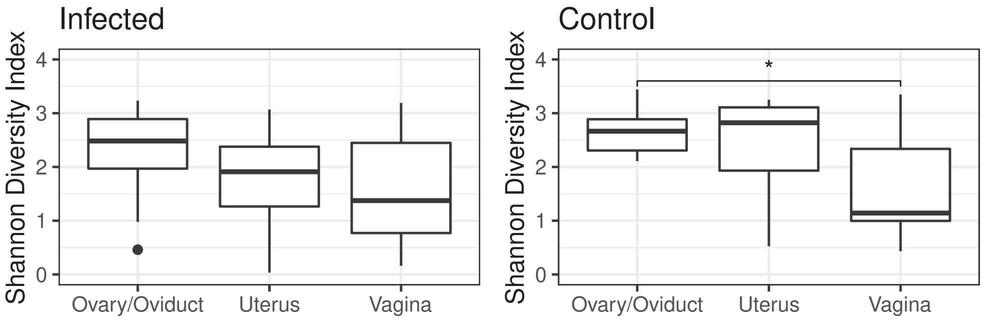


(a) Shannon Diversity Index

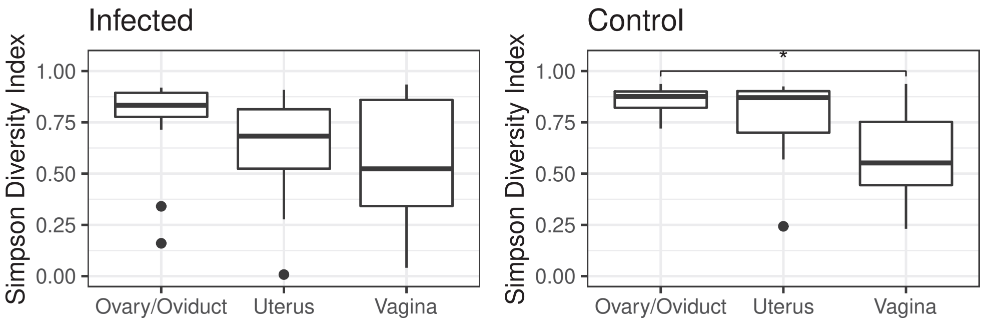


(b) Simpson Diversity Index

Figure C.1: Boxplots of Shannon (panel (a)) and Simpson Diversity Index (panel (b)) of samples collected from the genital tract regions four weeks post infection for infected group (ZT15_I and ZT3_I) and control group (ZT15_C and ZT3_C). Statistical significance is indicated above the brackets: $*, p \leqslant 0.05$.

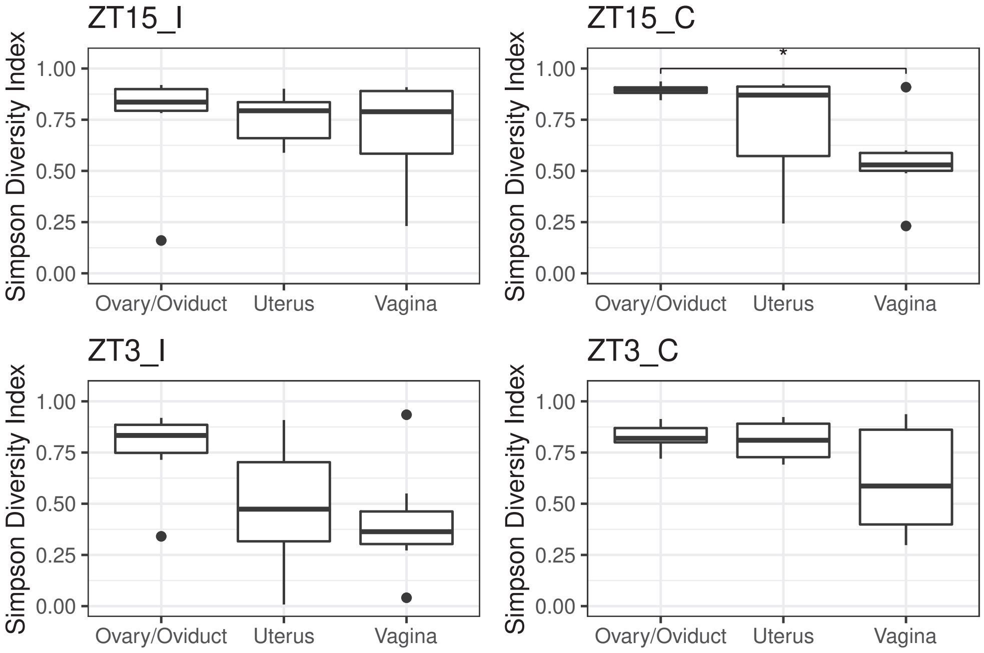


Figure C.2: Boxplots of Simpson Index (panel (b)) for samples collected four weeks post-infection by GTR for each group (ZT15_I, ZT15_C, ZT3_I, and ZT3_C). Statistical significance is indicated above the brackets: *, $p \leqslant 0.05$.

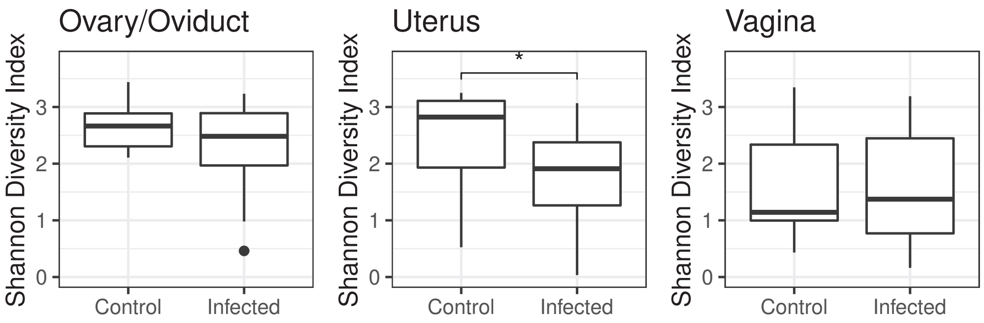


(a) Shannon Diversity Index

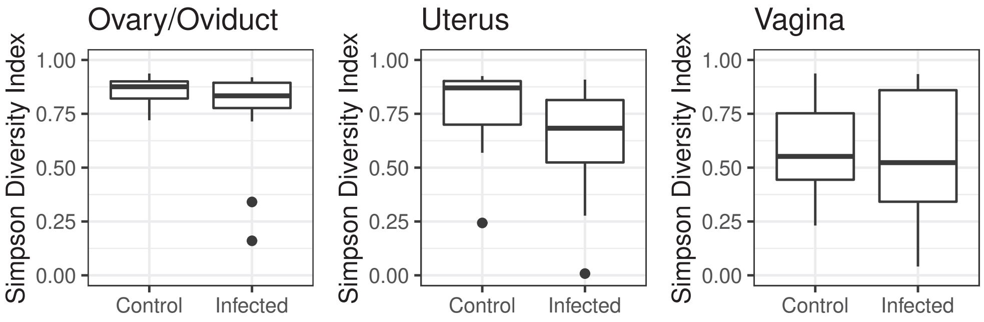


(b) Simpson Diversity Index

Figure C.3: Boxplots of Shannon (panel (a)) and Simpson Diversity Index (panel (b)) per genital tract region for samples collected from infected group (ZT15_I and ZT3_I) and control group (ZT15_C and ZT3_C) four weeks post infection. Statistical significance is indicated above the brackets: $*, p \leqslant 0.05$.

### D PCoA for Week 4 Samples

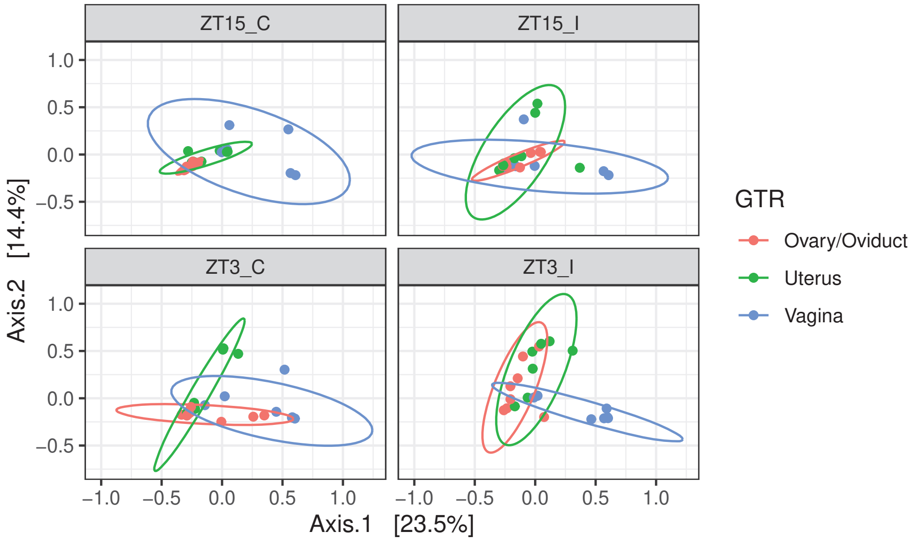


Figure D.1: PCoA results for beta diversity metrics among genital tract regions (GTRs) by treatment group, showing Bray-Curtis distance, for samples collected four weeks post infection. The ellipses for clustered samples assume multivariate t-distribution. The first axis explains 23.5% of the variability and the second axis explains 14.4% of the variability in the data of samples collected four weeks post infection.

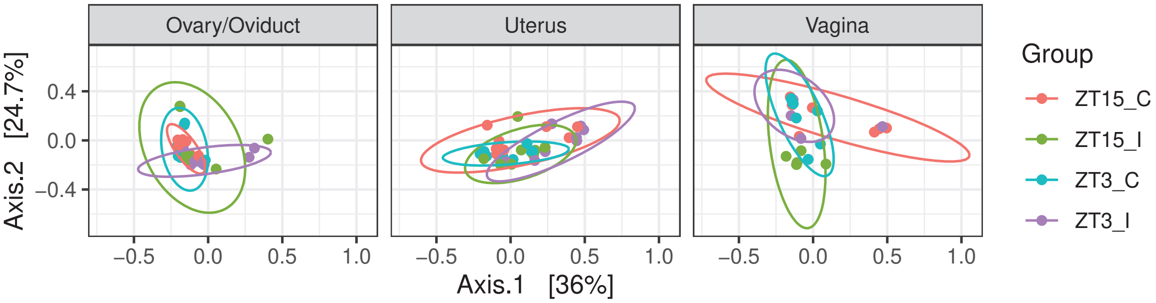


Figure D.2: PCoA results for beta diversity metrics among treatment groups by GTR, showing W-Unifrac distance, for samples collected four weeks post infection. The ellipses for clustered samples assume multivariate t-distribution. The first axis explains 36% of the variability and the second axis explains 24.7% of the variability in the data of samples collected four weeks post infection.

### E CCA for Week 4 Samples

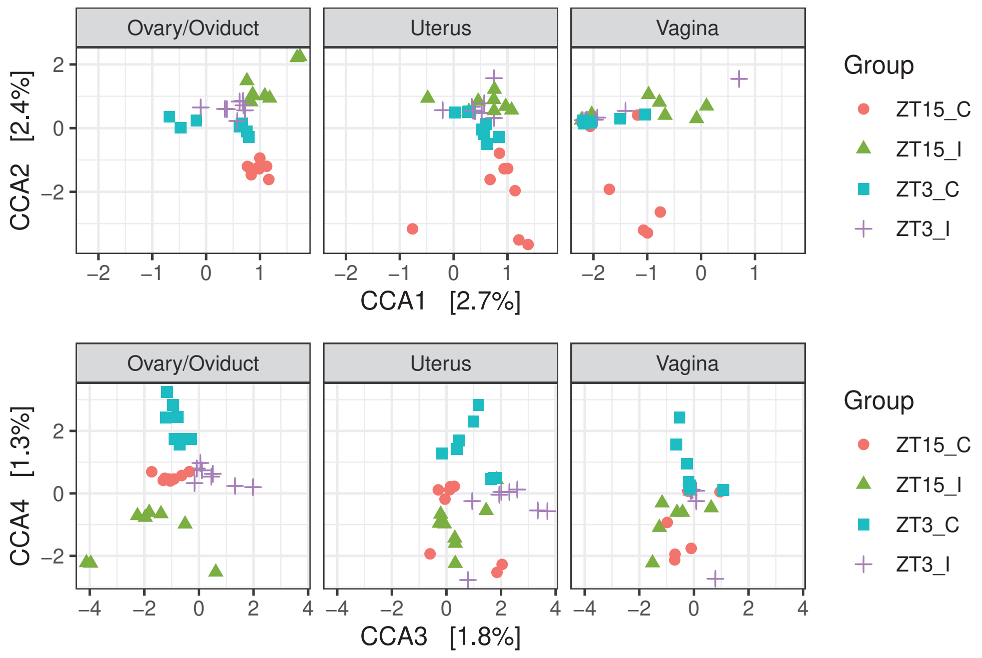


Figure E.1: CCA Ordination diagram with first four constrained axes for samples collected four weeks post infection, faceting by GTR. 2.7% , 2.4% , 1.8% and 1.3% of the constrained inertia in the data of samples collected four weeks post infection is explained by the first, second, third, and fourth constrained axis, respectively.
